# PANDAA-monium: Intentional violations of conventional qPCR design enables rapid, HIV-1 subtype-independent drug resistance SNP detection

**DOI:** 10.1101/795054

**Authors:** Iain J. MacLeod, Christopher F. Rowley, M. Essex

## Abstract

Global efforts to ensure that 90% of all HIV-infected people receiving antiretroviral therapy (ART) will be virally suppressed by 2020 could be crippled by increases in acquired and transmitted HIV drug resistance (HIVDR), which challenge ART efficacy. The long-term sustainability of ART treatment programs is contingent on effective HIVDR monitoring yet current Sanger sequencing genotypic resistance tests are inadequate for large-scale implementation in low- and middle-income countries (LMICs). A simple, rapid, affordable HIVDR diagnostic would radically improve the treatment paradigm in LMICs by facilitating informed clinical decision-making upon ART failure. Although point mutation assays can be broadly deployed in this context, the primary challenge arises from extensive sequence variation surrounding targeted drug resistance mutations (DRMs). Here, we systematically and intentionally violate the canonical principles of qPCR design to develop a novel assay, Pan-Degenerate Amplification and Adaptation (PANDAA), that mitigates the impact of DRM-proximal secondary polymorphisms on probe-based qPCR performance to enable subtype-independent, focused resistance genotyping. Using extremely degenerate primers with 3’ termini overlapping the probe-binding site, the HIV-1 genome is adapted through site-directed mutagenesis to replace secondary polymorphisms flanking the target DRM during the initial qPCR cycles. We show that PANDAA can quantify key HIV DRMs present at ≥5% and has diagnostic sensitivity and specificity of 96.9% and 97.5%, respectively, to detect DRMs associated with ART failure. PANDAA is an innovative solution for HIVDR genotyping and is an advancement in qPCR technology that could be applicable to any scenario where target-proximal genetic variability has been a roadblock in diagnostic development.

## INTRODUCTION

More than 23 million people living with HIV rely on the continued effectiveness of antiretroviral therapy (ART) to safeguard their health and survival *(1)*. The long-term success of treatment programs is contingent on effective monitoring of HIV drug resistance (HIVDR), which challenges ART efficacy and sustainability. ART should result in durable HIV replication suppression to sub-detection levels provided that ART adherence is uninterrupted, however, 10-30% of patients experience treatment failure annually (2–4). Without virological suppression, continuation of the same ART regimen leads to the emergence of acquired drug resistance (ADR), which limits the efficacy of current and future treatment regimens by rendering one or more of the antiretrovirals (ARVs) used in ART – or even whole drug classes – ineffective *(5)*. These virologically-unsuppressed individuals may transmit drug-resistant HIV to others, and individuals with transmitted drug resistance have more than double the odds of virological failure upon receiving ART *(6)*.

In low- and middle-income countries (LMICs), single treatment regimens are provided for first- and second-line ART *(7, 8)*, as volume procurement reduces the burden on healthcare infrastructures and facilitates programmatic expansion while reducing local clinical decision-making. HIVDR has increased as a of the global ART scale-up, particularly given the use of these proscribed ART regimens *(5)*. Although resistance genotyping allows clinicians to classify virological failure as resistance- or adherence-mediated, and to select an alternative ART regimen that confers the highest likelihood of virological success, in LMICs it is typically unobtainable until a patient fails to respond to two standardized regimens. A major premise for siloing resistance genotyping is that it is unactionable as a single standardized second-line regimen is available. Nevertheless, from an economic and healthcare-delivery perspective, the primary benefit is derived from preventing an ART switch in the 30-40% of non-drug-resistant patients with virologic failure due solely to adherence issues *(9–11)*. When virologic failure is adherence-mediated, second-line ART is unlikely to improve patient outcomes and prematurely restricts future treatment options.

Until recently, non-nucleoside reverse transcriptase inhibitors (NNRTIs) were the mainstay of first-line ART. The WHO now recommends dolutegravir due to its higher genetic barrier to resistance *(8)*. WHO guidelines recommend switching virologically suppressed patients from an NNRTI-based to a dolutegravir-based first-line regimen with the same nucleoside reverse transcriptase inhibitors (NRTI) backbone *(8)*. As such these patients retain their position in the ART regimen cascade on a first-line regimen. Those failing NNRTI-based ART will, on the presumption of resistance-mediated failure *(12)*, be switched to a standardized dolutegravir second-line regimen with an alternative NRTI backbone that includes zidovudine – a poorly-tolerated ARV – and a second NRTI with potential cross-resistance to the first-line NRTIs *(8, 13, 14)*. Continuing to exclude resistance testing from the treatment paradigm deprives those non-drug-resistant patients the opportunity to remain at the start of the ART cascade on a first-line regimen. For treatment programs, this will result in a substantially higher proportion of patients needing to access expensive third-line ART in the future.

Current resistance genotyping approaches are ill-suited for large-scale implementation in LMICs as they rely predominantly on Sanger sequencing, and more recently, next-generation sequencing (NGS). These approaches demand centralized expertise and resources, averaging 2–3 days from HIV RNA isolation to genotyping results and costing up to US$300 per patient *(11)*. Although genotypic resistance testing is recommended after second-line ART failure to optimize third-line ART, these constraints continue to prevent widespread implementation of this policy. As the number of patients failing first- and second-line ART increases, a straight-forward, high-throughput genotyping approach is needed to ensure the continued success of ART. Focused genotyping is a novel approach that exploits key genomic positions whereby the presence of a single DRM yields high predictive value of reduced ART efficacy and treatment failure. Focused genotyping of just six codons can detect major NRTI and NNRTI DRMs in >98% of patients who fail an NNRTI first-line regimen *(15)*. Inexpensive and high-throughput focused resistance genotyping could provide cost savings for national ART programs as treatment access continues to expand and the number of patients switching to alternative regimens continues to grow. The best candidate assay for focused genotyping are point mutation assays (PMAs) (e.g., allele-specific PCR, oligonucleotide ligation assay, and probe-based quantitative real-time PCR [qPCR]) *(16)*. PMAs have high detection sensitivities for single nucleotide polymorphisms (SNPs), however, are exquisitely sensitive to genetic variability near the SNP and return a false negative result in the presence of a secondary polymorphism (a proximal nucleotide change unrelated to drug resistance) *(17)*. Although several HIV-1 drug resistance PMAs have been developed for research use, DRM-proximal genomic variability has prohibited their broad clinical implementation for focused genotypic resistance testing *(18, 19)*. Even though its application for SNP detection in heterogeneous genomes has proven problematic *(20, 21)*, probe-based qPCR maintains the greatest potential for focused resistance genotyping in LMICs given its simple workflow, low cost and ubiquity in diagnostic labs across the globe. Despite decades of reagent development, there hasn’t been a major reconsideration of fundamental qPCR design principles since their inceptual codification *(22–24)*. To overcome the limitations of conventional qPCR, we systematically and intentionally violated canonical qPCR design principles to develop a novel PMA, Pan-Degenerate Amplification and Adaptation (PANDAA), that uniquely enables qPCR for universal, focused genotyping. PANDAA addresses high intra- and inter-patient HIV genomic variability by normalizing probe-binding regions. Through simultaneous mutagenesis during qPCR amplification, PANDAA replaces secondary polymorphisms that would otherwise inhibit probe hybridization and minimizes their impact on qPCR sensitivity and specificity. PANDAA is an innovative solution for HIVDR genotyping in LMICs and solves a problem that is currently addressed ineffectively by more complicated and esoteric techniques. More broadly, PANDAA is an advancement in qPCR technology that could be applicable to any scenario where target-proximal genetic variability has been a roadblock in diagnostic development.

## RESULTS

### Principles of PANDAA amplification and adaptation

We want to challenge the belief that HIV-1 genetic heterogeneity presents too high of a barrier for the widespread implementation of PMAs for HIV-1 drug resistance genotyping. We postulated that: 1) qPCR probe design guidelines do not accurately reflect advancements made in PCR reagent development; 2) short probes (<15 nt) with an melting temperature (*T_m_*) close to that of the primers can provide requisite sensitivity and specificity for focused genotyping assays; 3) a high degree of nucleotide degeneracy can be incorporated into primer design without negatively affecting qPCR performance; and 4) qPCR probe design should incorporate global or regional HIV subtype prevalence to account for the likelihood of encountering a given subtype within a random sampling of HIV-1 infected patients.

As the pivotal determinants of qPCR sensitivity and specificity, primer and probe design suitability are primarily determined by two interdependent factors: the oligonucleotide T_m_, governed principally by oligo length and GC content, and its complementarity with the target nucleic acid sequence. Sequence divergence within oligonucleotide binding sites are a significant source of assay error as primer-template mismatches lower T_m_ to weaken duplex stability with mismatches proximal to the primer 3’ terminus having the largest negative effect on qPCR performance. Longer oligonucleotides and/or higher GC content increase T_m_ to accommodate mismatch tolerance and offset instability, however, too high of a T_m_ will reduce assay specificity by introducing unintentional mismatch tolerance. Alternatively, primer-template mismatches may be mitigated using degenerate nucleotides although a balance must be achieved to minimize both amplification bias of heterogeneous quasi-species and non-specific amplification by off-target mispriming. Incorporating degenerate nucleotides into qPCR probe design, offers limited relief from secondary polymorphisms: a degenerate probe pool may contain variations with a T_m_ too low for adequate sensitivity, or conversely, too high to allow specific SNP discrimination. Furthermore, the inclusion of degenerate bases in probe-based qPCR to differentiate a drug resistance-conferring SNP flanked by variable regions cannot account for *de novo* polymorphisms that may arise in the probe-binding site nor address the underrepresentation of HIV-1 inter-subtype genomic diversity in sequence databases.

We intentionally violated the core qPCR oligonucleotide design principles to develop PANDAA (**Table S1**), a technique that mitigates the impact of DRM-proximal secondary polymorphisms on qPCR performance. Using extremely degenerate primers that overlap with the probe-binding site, the HIV-1 genome is adapted through site-directed mutagenesis during the initial qPCR cycles to replace secondary polymorphisms flanking the primary drug resistance SNP (**Fig. 1A–D**). This approach generates an amplicon population with a homogenous probe-binding site whereby the only point of nucleotide variation is at the DRM. PANDAA primers contain two distinct regions: a 3’ adaptor region (ADR) that is matched to the probe-binding site and a pan-degenerate region (PDR) that incorporates degenerate bases representative of nucleotide variability in the targeted primer-binding site upstream of the ADR. PANDAA primers include locked nucleic acids (LNAs), which act as molecular anchors to increase primer affinity for their target sequences and counter the thermodynamic instability of mismatches within the ADR (**Fig. 1E**).

**Fig. 1.**
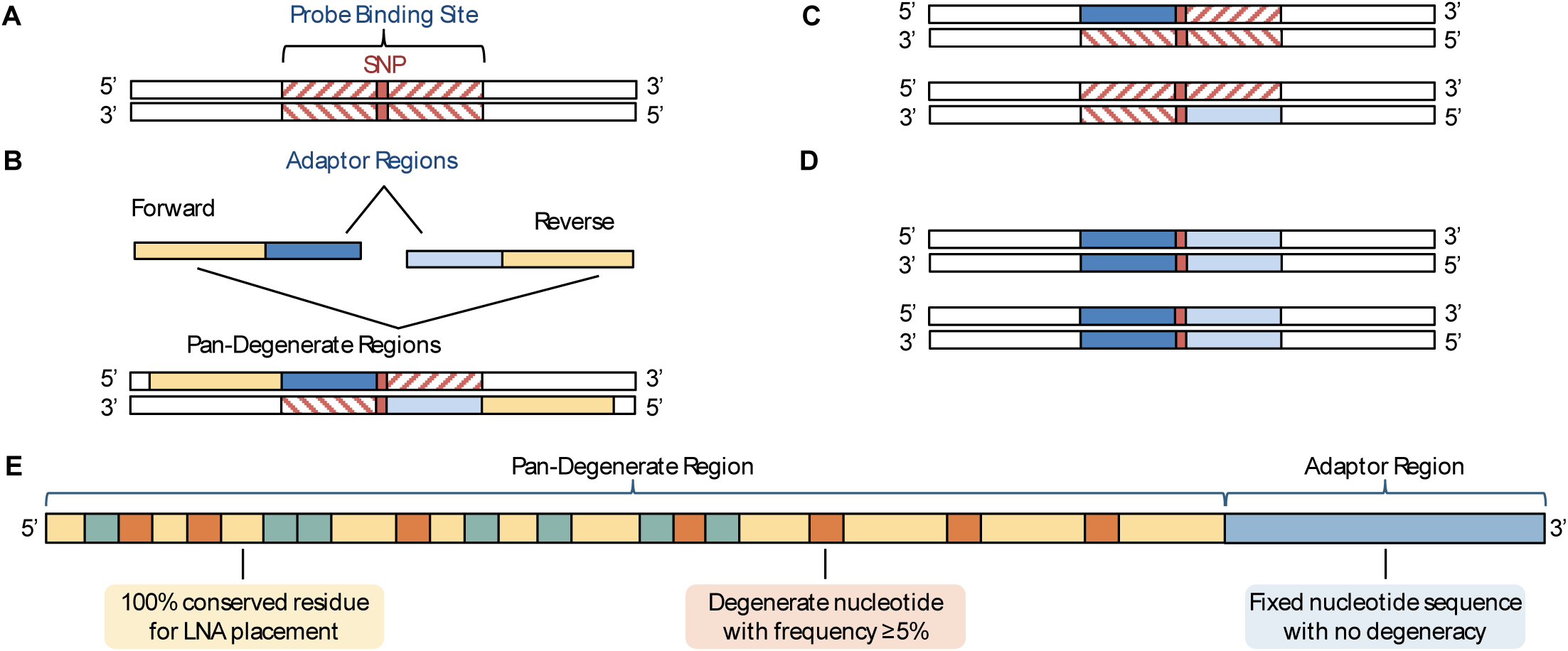
Overview of PANDAA method and primer design. (**A**) Heterogeneous genomes contain SNP-proximal secondary polymorphisms within the probe-binding site, preventing the use of probe-based qPCR for DRM discrimination. (**B**) PANDAA primers overlap with the probe-binding site, adapting the nucleotides proximal to the SNP of interest that would otherwise abrogate probe hybridization. (**C**) and (**D**) As qPCR proceeds, newly-generated amplicon will contain probe-binding sites flanking the SNP that are perfectly complementary to the probe. (**E**) PANDAA primers contain two key features: a 3’ ADR that is matched to the probe-binding site and a pan-degenerate (PDR) region that incorporates the nucleotide degeneracy observed in the primer-binding site of the target. The PDR is designed to account for the high degree of variability in primer-binding sites. LNA bases are incorporated into the primer 5’ region at 100% conserved positions to offset the thermodynamic instability of mismatches between the primer ADRs and the template.

### Probe-binding site variability around HIV-1 DRMs

The first step in the PANDAA design pipeline is to determine the probe sequence that confers specific DRM SNP discrimination. This subsequently instructs design of the PANDAA primer ADR for target adaptation. Using ~95,000 unique patient-derived HIV-1 sequences, the probe sequence is derived from the most prevalent probe-binding site allele (**Table S2**). Three methods have predominantly been used for HIV-1 oligonucleotide design, although not all of these methods account for inter-subtype diversity (**Table S3**). We developed an alternative approach that weights the frequency of a probe-binding site allele by adjusting for subtype prevalence (**Table S4**), which ensures an increased likelihood of encountering a matched target and mitigates bias from the unbalanced subtype distribution in publicly available sequence databases.

### Validation of PANDAA probe design

To provide mismatch tolerance during adaptation, PANDAA primers have a T_m_ 65–75°C. With conventional qPCR design, the probe T_m_ should be 8–10°C higher than the average primer T_m_, an ostensibly immutable constraint to ensure the probe out-competes the quick hybridization of the primers to their complementary strand during the annealing step *(22, 23)*. If we applied these canonical design rules, a PANDAA probe would have a T_m_ >75°C and would provide little to no DRM SNP discrimination. Therefore, we designed PANDAA probes with a T_m_ close to the 60°C assay annealing temperature, which favors the use of shorter probes therefore reducing the number of probe-binding site secondary polymorphisms to be adapted. We initially validated PANDAA to discriminate the wild-type amino acid, lysine (K), of codon 103 in HIV-1 RT from the DRM, arginine (N), arising from an A→C transversion at the third nucleotide, using a set of DNA templates incorporating 19 probe-binding site allele for codon 103 (**Table S2**). When experimentally validating *in silico* probe designs, we determined that PANDAA probe length could be reduced to as few as 12nt by incorporating stabilizing nucleotide modifications, such as the minor groove binding (MGB). DRM discrimination as a function of probe length was determined empirically given the inaccuracy of TaqMan-MGB probe T_m_ predictions (**Fig. 2A**). LNAs were found to have comparable performance to MGB probes although LNA nucleotide placement also had to be determined empirically (**Fig. 2B**) *(25)*.

**Fig. 2.**
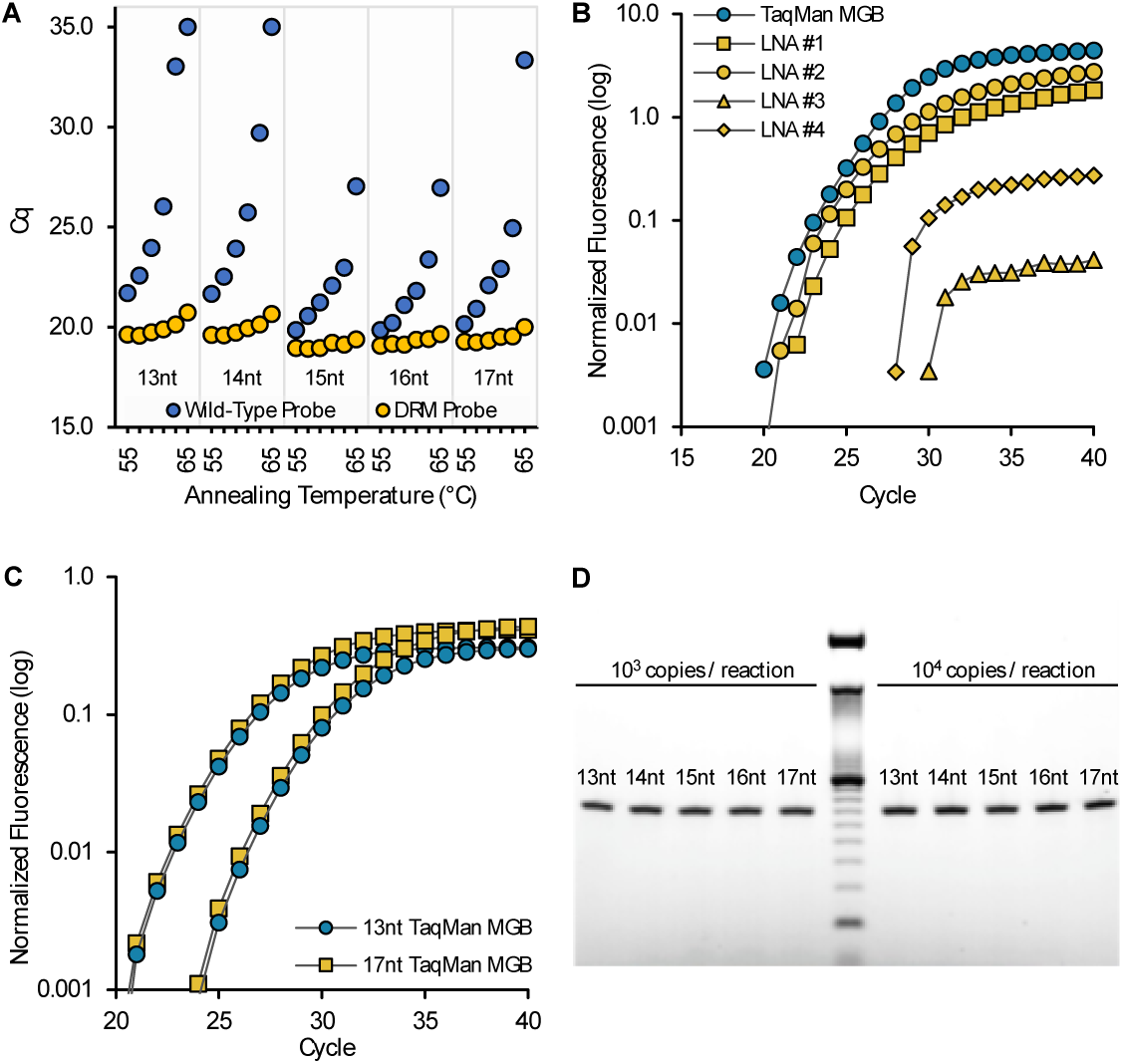
Optimization of PANDAA probe design. Discrimination of the K103N AAC DRM was evaluated using conventional qPCR with PANDAA primers lacking the 3’ ADR using template 001, which encodes the K103N DRM and does not contain additional secondary polymorphisms within the probe-binding site. (**A**) TaqMan probe hybridization properties differed from the predicted T_m_. 13–17 nt TaqMan-MGB probes were designed for 103 wild-type and DRM codons. Sensitivity and specificity were evaluated using an annealing temperature gradient from 55–65°C in 2°C increments. Although predicted T_m_ values ranged from 59–63°C, there was little impact sensitivity with the AAC probes (yellow circles) for any probe length as annealing temperature increased; however, non-specific hybridization of the AAA probe (blue circles) to the mismatched AAC template was reduced as annealing temperature increased. (**B**) The performance of four 5’ hydrolysis probes with various placement of LNA nucleotides (yellow) was compared to a TaqMan-MGB probe (blue) of the same length. (**C**) Increasing probe length did not reduce sensitivity, as 13-nt probes (blue circles) had similar Cq values to 17-nt probes (yellow) at 10^4^ and 10^3^ copies per reaction. (**D**) To confirm that amplification efficiency was not proportionally inhibited as the number of overlapping nucleotides increased, PANDAA reactions utilizing 13–17 nt probes were resolved on a 4% agarose gel, demonstrating comparable band intensities of a 66-bp amplicon across all probe lengths at both 10^3^ and 10^4^ DNA copies per reaction (middle lane: 10-bp DNA). Furthermore, no non-specific products were evident. Results represent median of six replicates.

The 3’ ADR of the forward and reverse PANDAA primers compete with the probe for the same amplicon binding site during probe-binding site adaptation. Longer probes may out-compete the primers for the same binding site and could reduce amplicon generation. Nevertheless, despite our disregard for the qPCR design principle prohibiting primer- and probe-binding site overlap, PANDAA neither reduced qPCR amplification sensitivity nor negatively impacted assay specificity. We evaluated 13–17 nt TaqMan-MGB probes targeting the K103N DRM codon with a DNA template containing no probe-binding site mismatches (template 001) and forward and reverse PANDAA primers with 6-nt ADRs. Despite binding-site competition between the primer ADRs and probes, PANDAA performance was equivalent across all probe lengths with a median Cq of 23.6 (IQR, 23.5–23.7) cycles at 10^4^ copies and 27.1 [interquartile range (IQR), 27.1–27.2] at 10^3^ copies (**Fig. 2C**). We confirmed that comparable qPCR performance was an artifact of the higher T_m_ and faster hybridization kinetics of longer probes masking a reduction in amplicon yield. Equivalent yields of a 66-bp amplicon were resolved by agarose gel electrophoresis, which also confirmed that complementarity between the probe and ADRs did not lead to the accumulation of non-specific products (**Fig. 2D**).

### Validation of PANDAA primer design

To mitigate the thermodynamic instability arising between primer ADR-template mismatches, the majority of a PANDAA primer is comprised of the PDR, which incorporates degenerate bases that reflect sequence diversity across the primer-binding site. We compared the performance of the non-degenerate consensus primers to the PANDAA primers 2830F and 2896R with degenerate bases included at positions with nucleotide variability ≥5% (95% consensus). Both primer sets contained the same 3’ ADR. Using template 001 diluted from 10^6^ to 10^2^ copies per reaction, the PDR increased PANDAA performance when compared to non-degenerate consensus primers (**Fig. 3A and B; Table S5**). We incorporated degenerate bases in the PDRs of 2830F and 2896R to represent the 95–99% consensus (i.e., nucleotides with a frequency from 1–5%) to determine empirically the assay tolerance for primer degeneracy between 672- and 19,968-fold (**Table S6**). Using template 014, which has probe-binding site mismatches in both the forward and reverse ADRs, PANDAA sensitivity improved when 2830F PDR degeneracy represented the 96/97% consensus: a ~15-fold increase was observed when combined with the 2896R-95% reverse primer and a ~22-fold increase with the 2896R-99% reverse primer (**Fig. 3C, Table S7**). Increasing 2830F degeneracy further did not improve sensitivity, and 2830F-99% exhibited no improvement in sensitivity relative to 2830F-95%. This reversal of amplification improvement was likely due to a reduction in effective primer concentration relative to primer degeneracy: 2830F-99% (18,432-fold) compared to 2830F-96/97% (1,536-fold) degeneracy (**Table S6**). Comparable SYBR green experiments resolved single amplicon peaks demonstrating that the reduced amplification efficiency of 2830F-99% did not arise from reaction component sequestration by non-specific product formation (**Fig. 3D, Fig. S2**). A similar pattern, albeit with a reduced relative increase in amplification, was observed when using the 001 template (**Table S7**). Additionally, co-expressed DRMs (i.e., those proximal to the primary DRM of interest) were incorporated as additional degenerate positions. For K103, this included codons 100 and 101 in primer 2830F and 106 in 2896R, all of which increased degeneracy without reducing performance (**Fig. 3E**).

**Fig. 3.**
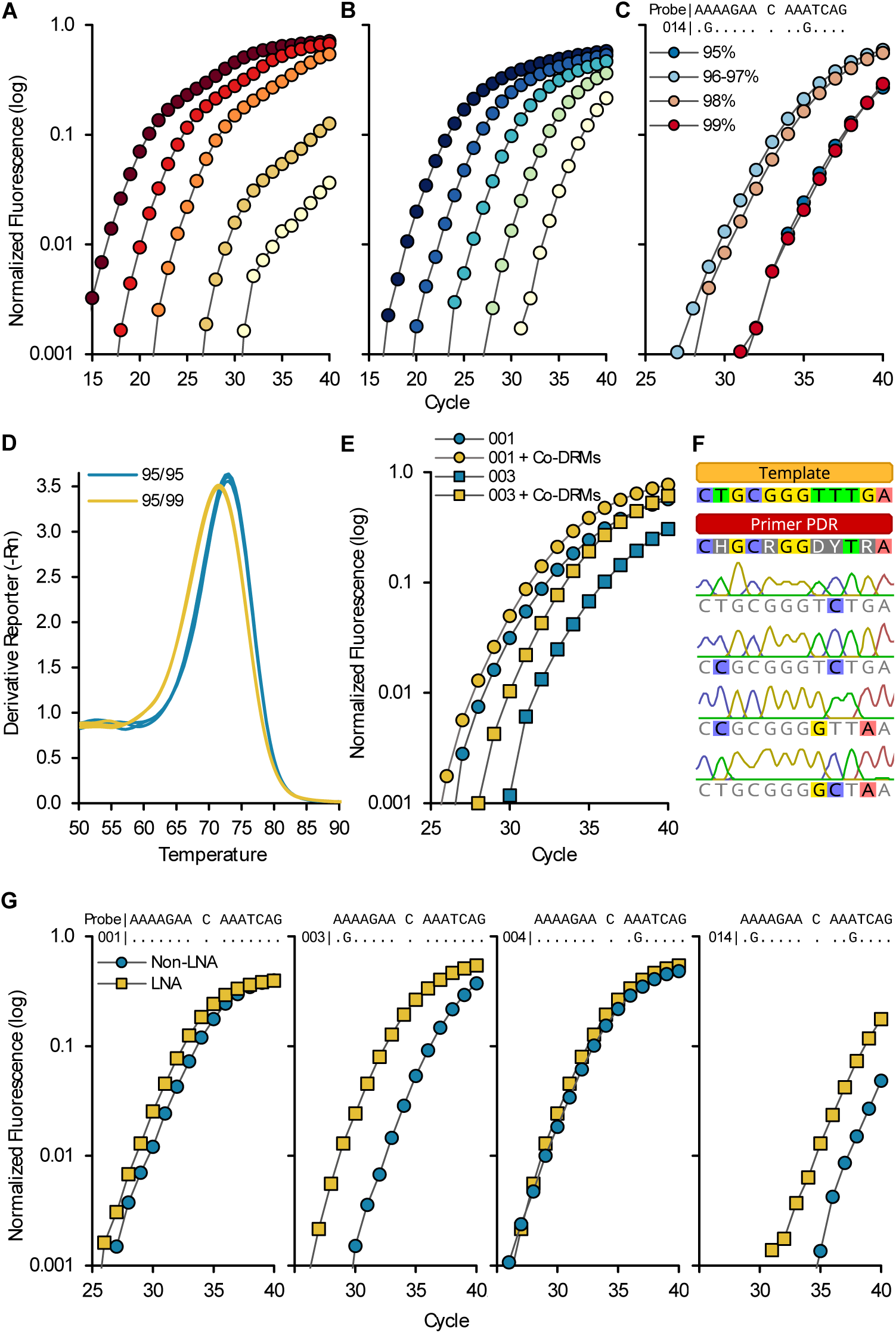
Optimization of PANDAA primer design. Consensus versus PANDAA primers. The 001 template was diluted from 10^6^ to 10^2^ copies per reaction and was amplified using either the majority consensus primers with no degenerate bases (**A**) or PANDAA primers (**B**). Both sets of primers contained the same 3’ ADR. A single mismatch was present in both the forward and reverse consensus primers (**Table S5**). Using the K103N DRM-specific probe, both primer sets showed similar sensitivity down to 10^4^ copies; however, the lowest copy number that could be accurately quantified using the consensus primers was 10^4^ copies compared to 100 copies for the PANDAA primers. Although <10^4^ copies were detectable using consensus primers, the copies were not accurately quantifiable. PDR degeneracy, co-expressed DRMs, and melt curve: (**C**) Using 10^3^ copies of template 014, which contains a mismatch in each ADR, optimal PDR degeneracy has to be determined empirically, as there was a tipping point after which increased degeneracy had no effect on PANDAA performance. (**D**) The 2830F-99% PDR variant was used with either the 2896R-95% (blue) or 2896R-99% (yellow) variant in a SYBR green qPCR with the 014 template. A similar relationship between PDR degeneracy and reaction efficiency to that with probe-based PANDAA was observed. The lower amplicon T_m_ and broader melt curve with the 2896R-99% primer reflected the wider range of predicted GC% content compared to 2896R-95% (25.8–58.1% versus 29.0–54.8%, respectively), and therefore the wider primer T_m_ range (63.0–78.7°C versus 64.2–77.3°C, respectively). (**E**) Additional degeneracy was introduced at co-expressed DRMs. Performance was evaluated using template 001 (circles) or template 003 (squares). Primers with co-expressed DRMs (yellow) were compared to the previous iteration of PANDAA primers (blue) and were found to improve sensitivity. (**F**) Single-clone sequencing after performing PANDAA on a homogeneous DNA template demonstrated that adaptation was occurring within the PDR at positions containing degenerate bases. (**G**) Inclusion of LNA nucleotides at 100% conserved positions in the PDR increased sensitivity in templates containing probe-binding site mismatches in the forward ADR, reverse ADR, or both.

PANDAA was shown to adapt the entire primer-binding region, demonstrating that there is a broad spectrum of degenerate primer utilization. Using two homogenous templates with different 2830F and 2896R primer-binding sites, single-clone sequencing showed that adaptation occurred within the PDR binding region at primer positions containing degenerate nucleotides (**Fig. 3F, Fig. S3**). As these sequences represent the predominate populations at the completion of the qPCR, those containing multiple substitutions may not represent adaptation to that sequence by a single degenerate primer. Rather, substitutions will have occurred in a stepwise manner, with one or two changes incorporated at a time. This approach increases the effective primer concentration with each cycle as progressively more primer variations in the degenerate pool can hybridize with newly adapted amplicon.

Another approach to compensate for primer T_m_ reductions arising from ADR-template mismatches is to increase the primer length. Adopting this approach would require a further increase in primer degeneracy due to the high genomic heterogeneity of HIV-1. As an alternative strategy, we offset the thermodynamic instability arising from primer 3’ ADR-target mismatches by incorporating LNA nucleotides at 100% conserved positions. This allows the PDR to counterbalance possible primer T_m_ reductions and was shown to further enhances PANDAA sensitivity (**Fig. 3G**). Together, these iterative refinements to the PANDAA primer design—the empirical determination of both optimal degeneracy and the placement of LNAs at preferred positions that are 100% conserved—culminate in an assay that not only is highly tolerant of primer-template mismatches but also can eliminate DRM-proximal secondary polymorphisms, which have been a constraint in conventional qPCR design for decades.

### Resolution of probe-binding site mismatches

To demonstrate the ability of PANDAA to resolve probe-binding site mismatches, we compared PANDAA to conventional qPCR with PANDAA primers lacking the 3’ ADR and using 10^6^ copies of DNA template for each of the 19 probe-binding site variants represented in **Table S2**. A high copy number was used to ensure that inefficient amplification by conventional qPCR was not unnecessarily constrained. PANDAA increased sensitivity for all templates regardless of the position or number of mismatches (**Table 1; Fig. 4A–C**). Where probe binding was completely inhibited by a secondary polymorphism using conventional qPCR (n = 7), probe binding and DRM detection were rescued by PANDAA to within a median of 2.3 cycles (IQR, 0.6 to 6.8) from the perfectly matched template 001 (**Table 1**). The difference in performance between conventional qPCR and PANDAA was unlikely due to differences in amplification efficiency; both conventional qPCR and PANDAA had similar median Cqs for template 001, which does not require adaptation. Thus, PANDAA can adapt one or more secondary polymorphisms in the K103 probe-binding site, independent of the mismatch position relative to the SNP and the type of mismatch. Single-clone sequencing of PANDAA amplicons verified that adaptation occurred in the probe-binding site (**Fig. 4D**). Furthermore, PANDAA was successful in a one-step RT-qPCR when using RNA, demonstrating that PANDAA primer design does not impede cDNA synthesis (**Fig. S4**).

**Table 1.**
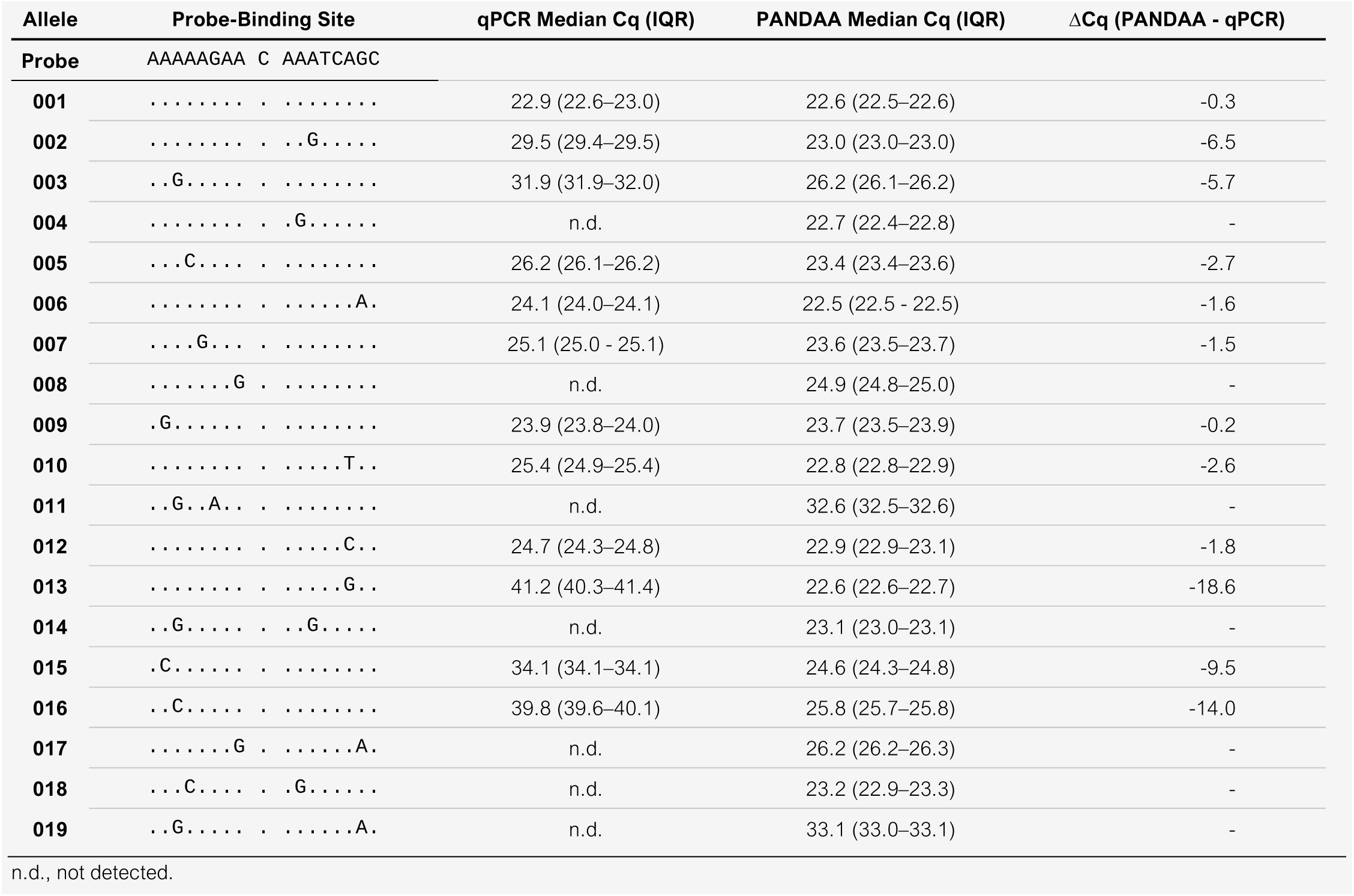
PANDAA improves sensitivity to detect single nucleotide changes in HIV-1 compared to conventional qPCR.

**Fig. 4.**
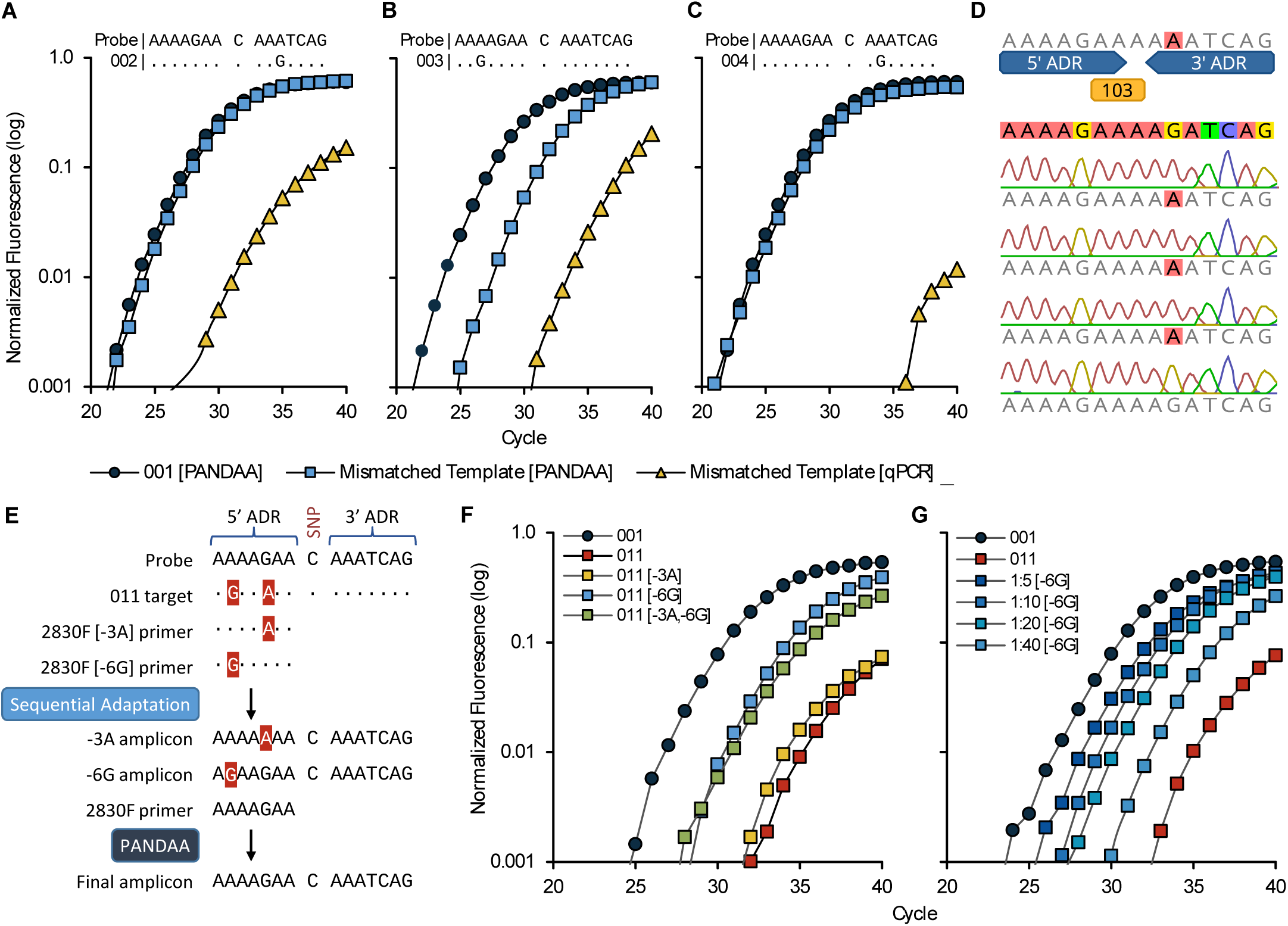
PANDAA adaptation of the probe-binding site rescues qPCR performance. With 10^4^ copies per reaction of DNA, conventional qPCR using PANDAA primers lacking the 3’ ADR was compared to PANDAA for the four most common probe-binding sites (001 to 004) flanking the 103 codon. Template 001 (blue circles), which does not contain additional secondary polymorphisms within the probe-binding site, was included as a reference in each experiment against which the target containing a secondary polymorphism was compared when using PANDAA (blue squares) versus conventional qPCR (yellow triangles). (**A**) PANDAA restored detection of template 002 to the same level of detection as a target with no secondary polymorphisms. (**B**) Detection of template 003 was increased almost 40-fold when using PANDAA compared to conventional qPCR. (**C**) Conventional qPCR did not detect template 004, whereas PANDAA restored detection close to that observed with template 001. Results are the representative median of six replicates. (**D**) Single-clone sequencing of the PANDAA amplicon demonstrated adaptation of the probe-binding site when using template 004 as the target. Adaptation was seen in 42/44 clones (95.5%). (**E**) Sequential adaptation overcomes multiple ADR mismatches. Template 011 contains two mismatches in the 5’ ADR, which must be adapted by the forward primer. By including a low concentration of 2830F primers that contain only one of the two mismatches, adaptation can be performed in a stepwise manner such that the 2830F PANDAA primer must only adapt one secondary polymorphism rather than two. (**F**) Two primers, 2830F [-3A] and 2830F [-6G], each of which contains only a single mismatch, were added to separate PANDAA reactions at 10% of the 2830F forward PANDAA primer concentration, and adaptation performance was evaluated with 10^3^ copies per reaction template 001 or 011 DNA. 2830F [-3A] resulted in a 0.9-cycle decrease compared to the 011 template with only the standard PANDAA primer. 2830F [-6G] reduced the Cq by 4.8 cycles, whereas both sequential adaptation primers together led to a 4.1-cycle decrease. (**G**) A dose response was evident with 2830F [-6G] sequential adaptation from 2.5% to 20% of the 2830F forward PANDAA primer concentration.

### Sequential adaptation of multiple ADR mismatches

In cases for which >1 secondary polymorphism must be adapted by the same primer ADR, we hypothesized that PANDAA performance could be improved by promoting adaptation in a sequential manner through the inclusion of low-concentration PANDAA primers containing each of the individual ADR mismatches. Template 011 has probe-binding site secondary polymorphisms at the -3 and -6 positions relative to the SNP; thus, the forward PANDAA primer 2830F contains two ADR mismatches (**Fig. 4E**). By including a low concentration of 2830F that contains either the -3 or the -6 mismatch, our hypothesis was that the first few qPCR cycles would generate a heterogeneous amplicon pool in which some proportion of the amplicons would be adapted to match the probe-binding site only at the -3 position and the remaining amplicons would be adapted only at the -6 position. Thus, the template pool would be expanded to contain amplicons that have a single mismatch with the 2830F PANDAA primer, allowing PANDAA complete adaptation more efficiently (**Fig. 4E**). To confirm this hypothesis, we used two separate primers to evaluate sequential adaptation of the 011 template: 2830F[-3A], which retained the -3 G:A (template:primer) mismatch while adapting the -6 A:G mismatch, and 2830F[-6G], which retained the -6 A:G mismatch while adapting the -3 G:A mismatch (**Fig. 4E**). Relative to no sequential adaptation on template 011, sensitivity was improved by including a 1:10 ratio of either the 2830F[-3A] primer (1.8-fold) or the 2830F[-6G] primer (19.7-fold) (**Fig. 4F**). Combining both sequential adaptation primers at an equimolar concentration was less effective at improving sensitivity compared to the -6G primer alone (13.5-fold). The -3 G:A mismatch was preferentially adapted, given that the 2830F[-6G] sequential adaptation primer performed better than the 2830F[-3A] primer; furthermore, a dose-dependent effect was evident with this sequential adaptation primer (**Fig. 4G**).

We found that inclusion of the sequential adaptation primer 2830F[-6G] increased the sensitivity of other probe-binding sites with the -6 G:A mismatch, with a negligible −1.5-fold reduction in amplification of the remaining templates (IQR, −1.1 to −1.6-fold) (**Table S8**). This pro-amplification (pro-amp) effect was surmised to arise from the partial decoupling of the amplification process from its dependence on adaptation during the initial qPCR cycles. By increasing the pool of unadapted template, pro-amp could offset the amplification penalty associated with adaptation, such that a higher proportion of newly adapted amplicons can be generated within the first 10 qPCR cycles (**Fig. S5**). We evaluated allele-specific pro-amp that used a low-concentration ADR-matched primers (eight 2830F 5’ ADR variants and six 2896R 3’ ADR variants), representing the 19 probe-binding site allele variations used in these studies. Sensitivity improved with individual pro-amp primers for each probe-binding site alleles, with a median delta Cq (ΔCq) of −1.0 cycle compared to that without pro-amp (IQR, −1.3 to −0.1) (**Table S9**). This modest increase in sensitivity was negated when the individual pro-amp primers were pooled, which led to a median ΔCq of 0.0 cycles (IQR, −0.5 to 0.2).

### Sensitivity, specificity, and selectivity of optimized PANDAA

We applied these design refinements to three additional HIV-1 DRMs—K65R, Y181C, and M184VI—to produce a highly-specific, focused genotyping resistance assay to NNRTI-based ART regimens. These PANDAA assays were validated using two sets of five synthetic DNA templates incorporating probe-binding site alleles for RT codons 65, 103, 106, 181, 184, and 190 to cover ≥95% of patients. One set contained the wild-type codon (Integrated A 001-005) and the second contained the DRM-conferring nucleotide substitutions for K65R, K103N, V106M, Y181C, M184VI, and G190A (Integrated B 001-005) (**Table S10**). Using differentially labeled TaqMan probes to discriminate wild-type DNA from the K103N DRM, we were able to quantify both targets across a linear range down to 5 copies (*r*^2^ = 0.998) (**Fig. 5**). Similar performance was observed for the other three DRMs. PANDAA selectivity (i.e., the detectable proportion of DRM on a wild-type background) was assessed using mixed ratios of Integrated A (wild-type) and B (DRM) 001-005 DNA templates down to a DRM proportion of 1%. Extensive specificity evaluations of all PANDAA assays using human genomic DNA indicated that PANDAA maintained high specificity despite the presence of highly complex, non-HIV nucleic acid (**Fig. 5D**).

**Fig. 5.**
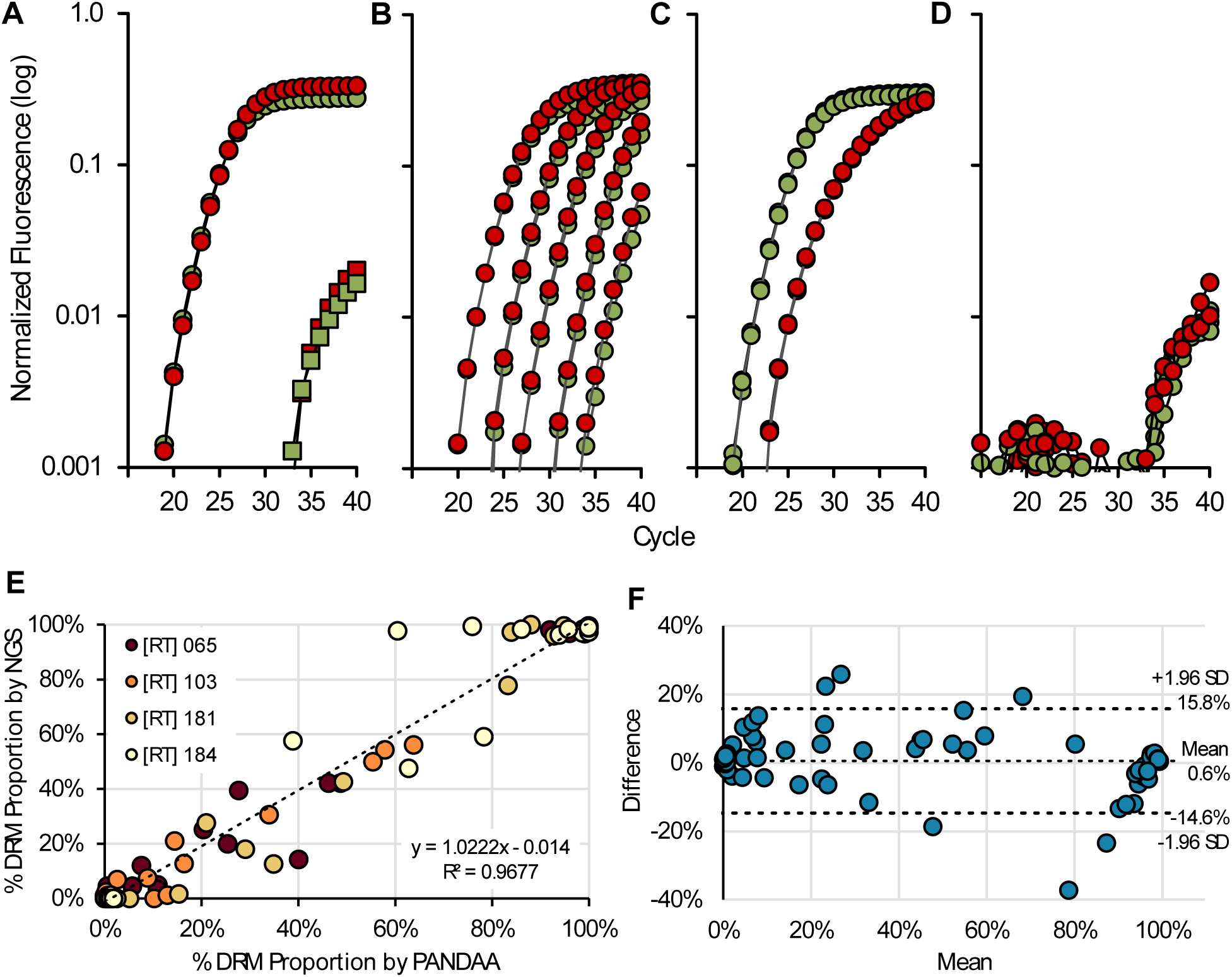
Validation of PANDAA performance. PANDAA was performed with differentially labeled TaqMan probes to discriminate wild-type DNA (VIC-labeled [green]) from the K103N DRM (FAM-labeled [red]). (**A**) Using either 100% wild-type DNA or 100% mutant 014 template DNA at 10^5^ copies per reaction, non-specific DRM signal (red squares) can clearly be differentiated from the specific wild-type signal (green circles) in wild-type only reactions. Similarly, the non-specific wild-type signal (green squares) can be distinguished from the specific K103 DRM signal (red circles). (**B**) PANDAA was performed on 10-fold dilutions of a 1:1 mixture of 10^5^ to 10 total DNA copies; thus, wild-type (green circles) and mutant DNA (red circles) were present at 50% of those quantities. (**C**) Mixed populations of wild-type (green) to mutant DNA (red), representing 10% K103N DRM at 10^5^ total copies of DNA. (**D**) Negative control using human genomic DNA. Results are representative of a minimum of six replicates of each dilution series. The x-axis represents the number of qPCR cycles and the y-axis represents the log normalized fluorescence. (**E**) Correlation of PANDAA with NGS. The proportions of K65, K103N, Y181C, and M184VI mutations were quantified by Illumina MiSeq NGS. Pearson’s correlation coefficient showed a strong agreement between the DRM proportions quantified by PANDAA and those quantified by NGS (r = 0.9837; 95% CI: 0.9759–0.9890; P < 0.0001) (**F**) A Bland-Altman plot of the agreement between DRM quantification by PANDAA and NGS shows a mean bias of 0.6% (±7.8%) with 95% limits of agreement (dotted lines) ranging from −14.6% to 15.8%.

### Comparison of PANDAA with resistance genotyping by population sequencing and NGS using clinical samples

We next evaluated PANDAA using 72 clinical samples from patients that experienced virological failure on an NNRTI-based first-line ART regimen. All samples were previously genotyped by population sequencing and probe-binding site mismatches were present in 18-43% of patients (**Table S11**). The associated PCR amplicons stored from this genotyping workflow were diluted prior to focused genotyping by PANDAA of RT codons 65, 103, 181, and 184. PANDAA showed excellent overall agreement with population sequencing, demonstrating 97.6% concordance (κ = 0.935; 95% CI: 0.887–0.983) (**Table 2**); 100% concordance was demonstrated with Y181C and M184VI. All three samples that were genotyped as K65R by PANDAA and as wild-type by population sequencing had ~5-9% electrophoretic mixtures when evaluated using Geneious. For the four discordant K103N results, the proportion of DRMs as determined by PANDAA was 11–15%, which is close to the cut-off used for population sequencing.

**Table 2.**
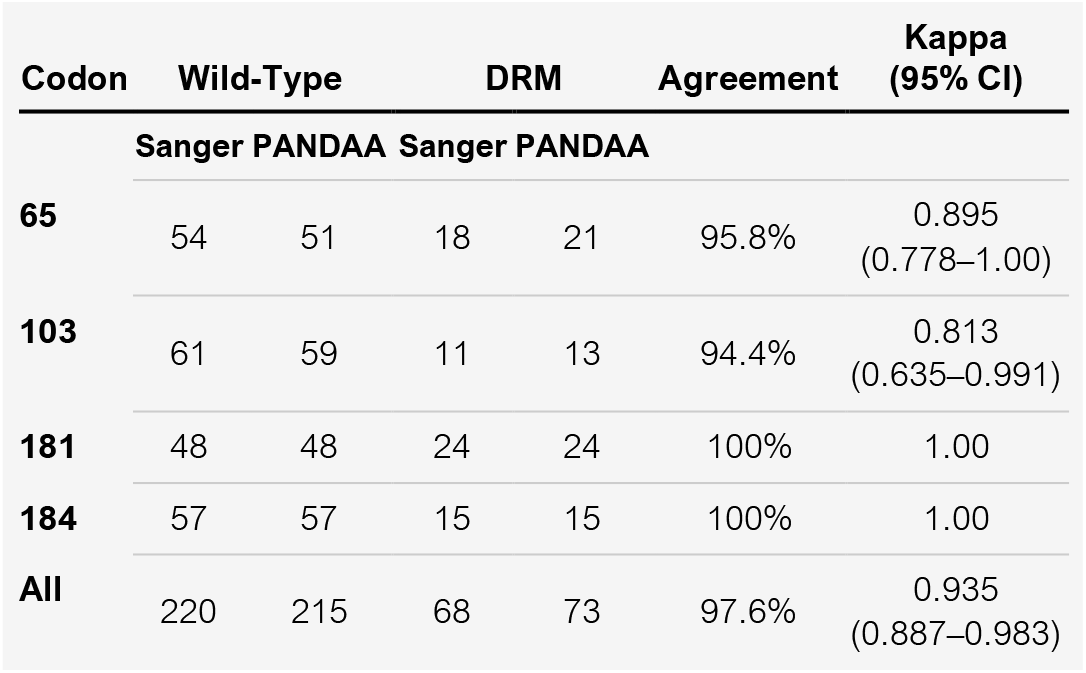
Agreement between PANDAA and population sequencing for four DRMs.

We evaluated diagnostic sensitivity to ascertain first-line ART failure, which was classified as the presence of ≥1 six failure-defining DRMs. Despite DRMs at two codon – V106MA and G190AS – not being covered by PANDAA in these studies, PANDAA had 96.9% sensitivity to accurately classify patients as treatment failures (**Table 3**). By drug class, PANDAA detected all patients with NRTI failure and 87.5% of those with NNRTI failure (**Table 3**). We analyzed a subset of 25 samples from the same cohorts using the NGS Illumina MiSeq platform to quantify DRM relative abundance and allow a comparison with the quantitative readout from PANDAA. Strong agreement was observed when PANDAA was compared to NGS for all four DRMs by Pearson’s correlation coefficient (*r* = 0.9837; 95% CI: 0.9759–0.9890; *P* < 0.0001) (**Fig. 5E, F**).

**Table 3.** Diagnostic sensitivity and specificity of PANDAA to determine first-line ART and drug class-specific failure.

|  | Sanger DRMs | Patient Coverage | PANDAA DRMs | Sensitivity (95% CI) | Specificity (95% CI) |
| --- | --- | --- | --- | --- | --- |
| **First-Line Failure** | K65R, K103N, V106M, Y181C, M184VI, G190AS | 99.2% | K65R, K103N, Y181C and M184VI | 96.9%<br>(84.3–99.5%) | 97.5%<br>(87.1–99.6%) |
| **NRTI Failure** | K65R and M184VI | 98.7% | K65R and M184VI | 100.0%<br>(85.7–100%) | 93.9%<br>(83.5–97.9%) |
| **NNRTI Failure** | K103N, V106AM, Y181C, and G190AS | 97.1% | K103N and Y181C | 87.5%<br>(71.9–95.0%) | 100.0%<br>(91.2–100%) |

## DISCUSSION

Without appropriate action, HIVDR will significantly undermine the global response to the HIV epidemic. Increasing pre-treatment resistance has exposed weaknesses in current HIV treatment algorithms – such as delays in viral load testing – that may encourage a higher-than-expected prevalence of treatment-emergent resistance despite the use of ARVs with a high genetic barrier to DRMs. Pre-treatment resistance testing can avert functional monotherapy in patients and will become a non-negotiable should two-drug regimens be implemented *(26)*. However, a uniform standard of care for all patients accessing treatment cannot be attained using existing genotyping diagnostics as they cannot withstand the resource and technical constraints of clinical and research laboratories in LMICs. To address these implementation shortcomings and the technical limitations of existing PMAs, we developed PANDAA by systematically violating the codified design principles of qPCR. PANDAA adapts the probe-binding site to mitigate the negative impact of sequence variability on qPCR performance thus enabling sensitive and specific DRM when conventional qPCR would have failed *(27, 28)*. PANDAA quantified DRMs regardless of the number or position of secondary polymorphisms, and we demonstrated the robustness of PANDAA adaptation by showing that >1 secondary polymorphism in the same ADR can be adapted sequentially by including a limiting concentration of single-mismatched primer. We show that PANDAA can quantify ≥5% DRM to return a focused genotyping result in ~90 minutes with RNA in a one-step RT-qPCR. We demonstrated excellent agreement between PANDAA and both population sequencing and NGS for four major RT DRMs. PANDAA has diagnostic sensitivity and specificity of 96.9% and 97.5%, respectively, in patients classified as first-line ART failures using conventional population sequencing genotypic resistance interpretation algorithms.

We established that three major qPCR design rules can be broken. First, we showed that primer- and probe-binding site overlap and sequence complementarity does not impede amplification or generate spurious non-specific products. Rules forbidding this are inherited from qPCR design employing longer probes and has not been empirically evaluated with MGB-stabilized 5’ hydrolysis probes until now. By minimizing probe length with TaqMan-MGB probes, we reduced the number of SNP-proximal polymorphisms to be adapted. Theoretically, primer-probe hybridization could generate a non-specific amplicon incorporating the complete probe-binding site. With PANDAA, complementary exists only with the primer ADR of the opposite orientation, e.g., the first 7 nt of the sense-oriented probe is complementary to the antisense primer 3’ ADR. Unfavorable hybridization of so few nucleotides at the 60°C annealing temperature, and the lack of a hydroxyl group at the probe 3’ terminus, minimize non-specific product formation.

Second, we showed that LNA bases increase tolerance for 3’ primer-template mismatches. Conventionally, any primer-template mismatch reduces thermal stability of the duplex. Those occurring within the last 4-5 nucleotides of the 3’ terminus disrupt the DNA polymerase active site and are the most detrimental to primer extension *(21, 29)*. We offset the thermal instability of 3’ mismatches in the PANDAAA ADR using LNAs to increase the T_m_ 2-8°C *(30)*. Other modified nucleosides, such as inosine, do not increase primer T_m_ and were excluded. Furthermore, DNA polymerase variants, which have varying levels of 3’ mismatch extension efficiency *(31, 32)*, are overlooked in design guidelines that prohibit 3’ mismatches. We leveraged the high rate of nonspecific nucleotide extension from 3’ mismatched bases of *Taq*, which is a critical design consideration for PANDAA *(31)*. With RNA templates, adaptation by the reverse primer ADR is delegated to the lower stringency cDNA synthesis step. Although amplification and adaptation efficiencies are interdependent, no primer-template mismatches are present in the newly synthesized amplicon once adaptation has occurred, allowing subsequent amplification rounds to proceed with increasing efficiency.

Lastly, we showed that PANDAA primers tolerate extreme degeneracy while ensuring low non-specific product formation. Increasing degeneracy is assumed to result in: 1) premature amplification plateau by proportionally lowering the concentration of unique target-matched primers able to prime amplification and will be consumed earlier in the reaction proportional reduction; 2) an excess of redundant primers that do not participate in amplification and promote non-specific product formation. Adaptation throughout the PDR-binding site by PANDAA results in low non-specific product formation; degenerate primers with PDR-template mismatches participate in productive amplification and is not limited to primers with perfectly complementary to the target. This facilitates an increasing proportion of the degenerate primer pool to participate in the reaction, further reducing the availability of degenerate primers to form non-specific products with the net effect of enhancing PANDAA reaction efficiency. Furthermore, we hypothesize that the high mismatch sensitivity of LNAs allowed degenerate PANDAA primers to impede non-specific product formation.

PANDAA differs from traditional degenerate oligonucleotide design by considering the complete primer- / probe-binding site as its own discrete genomic allele. Conventional design overestimates naturally occurring HIV genomic variation because each variant nucleotide is incorporated as a discrete change (**Table S3**). This generates oligonucleotides that do not occur naturally. Using the K65 codon as an example, the 95% consensus sequence for a 15-nt probe would generate 16 different probe sequences (**Table S3**). This is reduced to six sequences using our allele-based algorithm. Our approach reduces degeneracy further by using HIV-1 subtype prevalence to determine the probability of encountering a given subtype. By weighting primer- / probe-binding site allele frequency using subtype prevalence the likelihood of encountering a matched target is increased. In contrast, an uncorrected approach that determines the most frequent primer- / probe-binding site using an equal number of sequences from each subtype, introduces bias from low-prevalence subtypes, particularly circulating / unique recombinants.

Although shifting from a near absence of resistance genotyping for first-line failures in LMICs to point-of-care testing would represent a quantum leap in HIV care management, the work herein represents a significant advancement in diagnostic development. We aim to empower centralized laboratories with the ability to implement focused resistance genotyping as a reflexive diagnostic after a detectable viral load and PANDAA confers several advantages that support this goal. Unlike conventional qPCR, an a *priori* understanding of all probe-binding site variations does not need to be known; PANDAA can adapt *de novo* secondary polymorphisms within the probe-binding site. By virtue of the adaptation process, PANDAA is a subtype agnostic assay, an issue that has contributed to the high failure rate of commercial Sanger sequencing assays for non-B subtypes *(33, 34)*. As our algorithm can incorporate either global or regional subtype prevalence data, an assay that incorporates local HIV-1 sequence diversity in a specific geographical region can be readily designed, which may facilitate local research efforts in LMICs. Once PANDAA has been designed for optimal probe-binding site adaptation, there is substantial interchangeability between targeting a single or multiple DRMs at that codon without the need for primer redesign or re-optimization e.g., M184V or M184I/V. If there is no clinical utility in differentiating between DRMs, e.g., M184I/V, then probes can be labeled with the same fluorophore. This intrinsic flexibility removes redundant or superfluous detection reagents to maximize sample throughput and reduce costs. Furthermore, PANDAA’s inherent tolerance of extreme primer degeneracy favors the development of multiplexed assays to quantify DRMs at multiple codons in a single reaction, which we are currently investigating. The superior selectivity of PANDAA to detect of low-frequency DRMs, below the 15-20% threshold of Sanger *(35)*, is an additional strength that could further improve patient outcomes *(36, 37)*.

This study does have several limitations. Although PANDAA was optimized using synthetic templates designed from multiple HIV-1 subtypes (**Table S2**), we were only able to obtain samples from patients infected with HIV-1C. A direct comparison using prospectively collected samples from independent cohorts in multiple geographical regions is needed to evaluate the relative benefits and clinical utility of PANDAA compared to existing genotyping methods. This iteration of PANDAA was optimized for four DRMs conferring resistance to NNRTI-based regimens implemented in LMICs. Ongoing studies are expanding PANDAA to two additional first-line DRMs and for DRMs conferring resistance to second-line protease inhibitors. Although there are inherent limitations to the assumptions that we can make with regard to utilizing PANDAA for other highly polymorphic pathogens, we believe that the development of PANDAA as a multiplexed assay and its independent validation in a resource-limited setting will establish PANDAA as a platform diagnostic technology.

## MATERIALS AND METHODS

### Study Design

The objective of this study was to develop a rapid genotyping assay, PANDAA, for HIV-1 drug resistance mutations using quantitative real-time PCR. This required a in silico analysis of primer- and probe-binding site allele frequencies across all HIV-1 subtypes using a novel approach to weight allele frequency based on subtype global distribution. DRM-discriminating TaqMan MGB probes of various lengths were designed using the most common probe-binding site allele and the optimal length was determined empirically using synthetic DNA and RNA templates representative of DRM-proximal sequence variation. Degeneracy was incorporated within the primer pan-degenerate region to ensure ≥95% coverage of primer-binding site alleles and optimal degeneracy was determined empirically. The inclusion of thermostabilizing nucleotide modifications was assessed to improve PANDAA sensitivity and specificity. Optimized PANDAA primer and probe designs were compared to conventional qPCR to quantify the improved sensitivity of PANDAA in the presence of 19 probe-binding site containing mismatches at various positions relative to the DRM. Further evaluations of the limit of detection for PANDAA to quantify four DRMs was performed using synthetic DNA templates that represented ≥95% of probe-binding site alleles. Finally, PANDAA was compared to population and next generation sequencing using deidentified PCR amplicons derived from previous genotyping workflows from patients failing a first-line NNRTI-based ART regimen.

### HIV-1 Sequence Alignment

We searched the Los Alamos HIV public database (http://www.hiv.lanl.gov) for sequences within the genomic region 2550→3501 (HXB2 coordinates) from all subtypes – including recombinants – with a minimum fragment length of 500 nt. We selected a single sequence per patient, resulting in 93,611 sequences at the time of this study. Subtyping was determined by sequence-associated information from the Los Alamos database. Multiple sequence alignments were constructed using MAFFT 7.38 *(38)* in Geneious 11.1 (http://www.geneious.com) and checked manually. HIV-1 RT DRMs within the alignment were determined using the Stanford University HIV Drug Resistance Database (http://www.hivdb.stanford.edu) and reverted to the wild-type codon sequence *(39)*.

### Determination of Primer- and Probe-Binding Site Allele Frequencies

Alignments were analyzed using custom resequencing software written in Visual Basic (v7.1, Microsoft). The target region was extracted from each sequence and arrayed by subtype: A, 01_AE, 02_AG, B, C, D, F, and G. All other subtypes, including CRFs and URFs, were grouped as “Other”. Sequences with deletions or ambiguous nucleotides were excluded. Unique target sequences within each subtype array were identified and their prevalence determined. Any unique sequence with a prevalence <0.5% was excluded as potential sequencing error. The intra-subtype allele frequency (f_allele_) is then adjusted based on the subtype prevalence (*p_subtype_*): f_allele_ x p_subtype_. The final weighted prevalence of each target region allele is its cumulative frequency across all subtypes.

### Probe Design

For probes with an odd number of nucleotides and a single, centered DRM SNP, the upstream and downstream regions within the probe-binding site are of equal length: 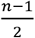 nucleotides where *n* is the probe length. To ensure that the discriminating SNP is biased toward the hydrolysis probe 3’ terminus, sense-oriented probes with an even number of nucleotides, have an upstream region 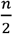 nucleotides, and downstream region 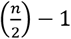 nucleotides. For antisense probes, the region lengths are swapped. Probes to detect the DRM were labelled with a FAM fluorophore and those for wild type with VIC.

### PANDAA Primer Design

A minimum of six PDRs of 30-40 nts were chosen for each forward and reverse primer-binding site. The 5’ terminal nucleotide would be placed at, or adjacent to, a conserved position at which an LNA nucleotide was incorporated. Two additional LNAs were placed downstream of 5’ terminus at 100% conserved positions based on previously reported design considerations *(30, 40, 41)*. The 95-99% consensus sequence was determined from the primer-binding site alleles with a cumulative frequency ≥95%. Primer ADR sequences were incorporated to represent the upstream and downstream regions of the optimal probe described above. Balanced ADRs are those for probe-binding sites of an odd-numbered length such that both PANDAA primer ADRs will be 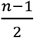 nucleotides. Final primer T_m_ predictions were calculated using Oligo Analyzer Version 3.1 *(42)* and a minimum of 36 pairwise primer combinations were empirically evaluated for optimal LNA placement and PDR degeneracy.

### PANDAA

PANDAA was performed using an ABI 7900 (Applied Biosystems). Briefly, 10μL reaction contained 5μL reaction buffer (Kapa Probe Fast, Kapa Biosystems), and optimized concentrations of forward and reverse PANDAA primers, VIC-labelled wild-type, and DRM-specific FAM-labelled probes. PANDAA reactions were incubated at 95°C for 3 min, followed by 10 three-step adaptation cycles of 95°C for 3 s, 50°C for 60 s, and 60°C for 30 s then 35 two-step amplification cycles of 95°C for 3 s, and 60°C for 90 s during which fluorescence data were captured. Reactions using RNA templates contained 15U MMLV reverse transcriptase (NEB), with an additional incubation step of 42°C for 15 min. SYBR qPCR was performed under the same conditions in the absence of PANDAA probes using Kapa SYBR Fast (Kapa Biosystems) with a melt curve stage included in the qPCR cycling protocol. Technical replicate number depended on the final copies / reaction: ≥10^4^ (n=4); ≥10^3^ (n=8); <10^3^ (n=12). Human genomic DNA at 0.05ng / reaction (Promega) was included as non-target nucleic acid negative control (n=8 replicates). Amplicons were resolved on 4% agarose EX e-gels (Thermo Fisher).

Raw qPCR fluorescence data were exported from Applied Biosystems SDS software. Background correction was performed using LinRegPCR *(43)*. For each target codon, PANDAA reaction efficiency was determined from standard curves of 1:1 mix of wild-type:DRM template across a dynamic range and calculated as efficiency (*E*) = 10 ^-1 / slope^ − 1. The quantification threshold (*N_q_*) was set at 0.05, which intersected with the exponential phase of the amplification curve for all targets at all copy numbers. This was used to determine the quantification cycle (*C_q_*) (the fractional number of cycles needed to reach *N_q_*). *C_q_* values were corrected for differences in probe-binding efficiencies to avoid biasing DRM proportion quantification due to asymmetric probe hybridization kinetics. The complete methodology for PANDAA data analyses can be viewed in the Supplementary Methods.

### Linearity, Sensitivity, Specificity and Selectivity

DNA concentration was determined by optical density, and copy number was calculated using the molecular weight of the nucleic acid before diluting to fixed copy numbers. Wild-type and DRM templates were mixed at a 1:1 ratio to a total of 10^6^ copies / μL and serially diluted two-fold to 8 copies / μL to determine PANDAA linearity for each target as well as limit of detection (LoD). Mixed ratios to provide a final DRM proportion of 25%, 10%, 5%, 2.5% and 1% were prepared in the same manner. The estimated LoD was determined by the lowest copy number whereby 95% of the replicates are positive and can be distinguished from the negative.

### Resistance Genotyping of Patient Samples

This study used de-identified PCR amplicon from the *Bomolemo* study, an observational cohort designed to demonstrate the tolerability and virological response to a fixed-dose efavirenz/tenofovir/emtricitabine ART regimen *(44)*. This study was conducted in Gaborone, Botswana between November 2008 and July 2011 by the Botswana-Harvard AIDS Institute and the Botswana Ministry of Health from whom Institutional Review Board approval was received. Viral RNA was isolated from patient plasma samples at virological failure and genotyped by population sequencing, as previously described *(45)*. Population sequencing chromatograms were analyzed using the automated resistance genotyping platform ReCall, with nucleotide mixtures called when the electropherogram peak was ≥10%. All patient-derived amplicon samples were diluted 1:1000 in dH_2_O supplemented with carrier tRNA from which 2μL was used in a PANDAA reaction, performed in triplicate by two operators.

NGS was performed using the MiSeq system (Illumina) with coverage to detect HIV-1 variants in 1% of the virus population. MiSeq libraries were prepared using the patient-derived amplicon with the MiSeq sequencing run performed at the Harvard Biopolymers Facility. Sequence quality was assessed using FastQC and QTrim was used to remove Illumina adapter sequences, reads below 36 bases, and leading/trailing low quality or N bases. Paired end reads were assembled into an HIV-1 Group-M consensus reference using Geneious Read Mapper software and non-synonymous detected using automated SNP calling. Results were verified using PASeq v1.4 (https://www.paseq.org) *(46)*.

### Statistical Analyses

Agreement between genotyping methods was determined by Pearson’s correlation coefficient. Kappa is a measure of the degree of non-random agreement between observers or measurements of the same categorical variable. Agreement is considered as good if kappa is between 0.60 and 0.80, and very good if greater than 0.80.

## Author contributions

I.J.M. and C.F.R. developed PANDAA and designed the performance experiments. I.J.M. was primarily responsible for data acquisition. C.F.R. fabricated the molecular clones. M.E. provided supervisory support. I.J.M., C.F.R., and M.E. discussed and interpreted the results, and wrote and edited the manuscript.

## Data and materials availability

PANDAA reagents (combined primers and probes) may be available through a material transfer agreement with the President and Fellows of Harvard College, or Aldatu Biosciences, Inc.

