## Supplementary Data for "PANDAA-monium: Intentional violations of conventional qPCR design enables rapid, HIV-1 subtype-independent drug resistance SNP detection"

### METHODS

#### Oligonucleotides

LNA-modified 5’ hydrolysis probes were synthesized by IDT, and TaqMan-MGB probes by Thermo Fisher. LNA-modified primers were synthesized from Exiqon whereas unmodified primers by from IDT.

#### Synthetic DNA

Synthetic double-stranded DNA was designed to evaluate the sensitivity and specificity of PANDAA. The 5’ region contains the T7 RNA polymerase promoter. Immediately downstream of the T7 promoter, and at the 3’ terminus, we included optimized primer-binding sites for SYBR green confirmation and normalization of copy number across different templates. Lyophilized DNA (geneStrings, Life Technologies) were resuspended in TE buffer to obtain a template master stock to 5ng /µL, which was quantified by fluorometry to provide an accurate stock concentration (Qubit dsDNA HS Assay Kit, Thermo Fisher). Templates were subsequently diluted in $dH_2O$ supplemented with carrier tRNA from *Saccharomyces cerevisiae* at 0.05 µg/µL (Sigma Aldrich) to provide a dilution series from $10^6$ copies/µL to 5 copies/µL.

#### Synthetic RNA

*In vitro* transcription of single-stranded RNA used 25ng synthetic DNA with the HiScribe T7 High Yield RNA Synthesis Kit (NEB). DNA template was removed from the RNA prep using RQ1 RNase-Free DNase and subsequently purified using RNeasy MiniElute Cleanup Kit (Qiagen; Hilden, Germany) with additional on-column DNA digestion. RNA was quantified by fluorometry to provide an accurate stock concentration (Qubit RNA HS Assay Kit, Thermo Fisher) and subsequently diluted in $dH_2O$ supplemented with carrier tRNA to $10^6$ copies/µL. Serial dilutions were performed in the same manner as those for synthetic DNA templates.

#### Single Amplicon Cloning and Sequencing

qPCR reactions were ExoSAP treated and purified (Wizard SV Gel and PCR Clean-Up System, Promega) prior to be ligated into the pMini vector (NEB PCR Cloning Kit, NEB). The ligation reaction was transformed into 10-beta Competent *E. coli* (NEB) and plated onto agar containing 100 µg/mL ampicillin. After overnight incubation at 37°C, inserts were screened by colony PCR using the Cloning Analysis Forward and Reverse primers (NEB) to identify a minimum of 45 single amplicon clones, which were then sequenced using the same screening primers (Genewiz). By sequencing 45 clones we had 95% confidence that we would detect sequence variants present in the amplicon population with a frequency of ≥10%: with *n* single amplicons sequenced, the probability (*P*) of missing a variant after screening *n* genomes is calculated as $f = 1 - \left(1 - P^{\frac{1}{n}}\right)$ when the variant comprises a fraction *f* (or less) of the virus population *(47)*.

#### PANDAA Data Analyses

Normalizes probe-binding efficiencies to avoid bias due to asymmetric probe hybridization kinetics. As the efficiency of PANDAA primers to amplify a target region is independent of whether the codon of interest encodes a wild-type amino acid or DRM then any differences in qPCR efficiency determined to exist between wild-type and DRM detection must arise due to differences in probe-binding efficiencies arising from small $T_m$ variances due sequence differences at the target SNP. The efficiency correction factor ($E_{correction}$) is determined by Equation 1 *(48, 49)*. The Cq of the DRM probe ($Cq_{(DRM)}$) is adjusted ($Cq_{(DRM)corrected}$) (Equation 2) to that which would have been obtained had both probes had the same hybridization efficiencies.

Although the wild-type and DRM amplification curves become parallel after efficiency correction, there will still exist difference in target Cq despite target nucleic acid being present in equal proportions for both probes. This second adjustment factor arises due to difference in probe fluorophore characteristics (i.e., background fluorescence, signal-to-noise ratio) *(49, 50)*. As target efficiencies have been already corrected $Cq_{shift}$ can be readily determined (Equation 3). Provided that $Cq_{shift}$ is constant across a dynamic range of input copy numbers at an equal ratio of wild-type and DRM templates then the final, adjusted DRM Cq is determined by equation 4 *(49, 50)*. This allows the proportion of DRM-harboring virus to be calculated using the common efficiency (equation 5).

**Equation 1**

$$E_{correction} = \frac{\log(E_{DRM})}{\log(E_{WT})}$$

**Equation 2**

$$Cq_{(DRM)corrected} = Cq_{(DRM)} \times E_{correction}$$

**Equation 3**

$$Cq_{shift} = Cq_{(DRM)corrected} - Cq_{(WT)}$$

**Equation 4**

$$Cq_{(DRM)shift} = Cq_{(DRM)corrected} - Med\left(Cq_{shift}\right)$$

**Equation 5**

$$Ratio = E_{WT}^{-\left(Cq_{(DRM)corrected} - Cq_{(WT)}\right)}$$

**Table S1. Comparison of canonical qPCR design criteria compared to those of PANDAA.**

| | Primers | | Probes | |
|---|---|---|---|---|
| **Design Condition** | **Conventional qPCR** | **PANDAA** | **Conventional qPCR** | **PANDAA** |
| **Oligonucleotide $T_m$** | 50–60°C | 65–75°C | 68–70°C<br>8–10°C > primer $T_m$ | 55–60°C<br>< primer $T_m$ |
| **GC content** | 30–80% | 30–80% | 30–80% | 30–80% |
| **Length** | 15–30 nt | 30–40 nt | 15–30 nt | 12–15 nt |
| **Distance between primer and probe** | 50 nt | Overlap | 50 nt | Overlap |
| **3′ terminus** | ≤2 GC in the last 5 bp<br>No mismatches | Mismatches tolerated | - | - |
| **Degree of nucleotide degeneracy** | Avoid | Up to 40,000-fold * | Avoid | Permissible |
| **Amplicon length** | 50–150 bp | 60–90 bp | - | - |

* not an absolute limit and should be determined empirically.

**Table S2. Frequency of probe-binding site polymorphisms around codon 103 of HIV-1 reverse transcriptase.**

| Allele | Prevalence | Cumulative | Accession | 15-nt Probe-Binding Site | Country | Subtype | RT Mutations |
|---|---|---|---|---|---|---|---|
| **Naturally Occurring Polymorphisms** | | | | | | | |
| **001** | 84.8% | 84.8% | AF443074 | AAAAGAA A/C AAATCAG | Botswana | C | |
| **002** | 4.2% | 88.9% | EF550533 | ....... . ..G.... | Ghana | CRF02_AG | |
| **003** | 4.2% | 93.2% | HM025038 | .G..... . ....... | Brazil | B | M41L, L100I |
| **004** | 1.9% | 95.0% | AF361877 | ....... . .G..... | Tanzania | C | V90I |
| **005** | 1.9% | 96.9% | GQ371960 | ..C.... . ....... | USA | B | |
| **007** | 0.8% | 97.7% | KC204777 | ...G... . ....... | Botswana | C | |
| **010** | 0.4% | 98.1% | EF394236 | ....... . .....T. | China | B | |
| **011** | 0.4% | 98.5% | JX447900 | .G..A.. . ....... | Thailand | CRF01_AE | |
| **012** | 0.2% | 98.7% | GU345235 | ....... . .....C. | China | B | |
| **013** | 0.2% | 98.9% | KC238186 | ....... . .....G. | Australia | B | D67N, T69D, K70R |
| **014** | 0.2% | 99.1% | JN000043 | .G..... . ..G.... | Cuba | CRF18_cpx | |

| Allele | Prevalence | Cumulative | Accession | 15-nt Probe-Binding Site | Country | Subtype | RT Mutations |
|---|---|---|---|---|---|---|---|
| **Co-Expressed DRMs in Probe-Binding Site** | | | | | | | |
| **006** | - | - | KJ176492 | ....... . ......A | South Africa | C | V106M |
| **015** | - | - | FJ688264 | C...... . ....... | Cameroon | A | D67E, V75M, K101P |
| **008** | - | - | KC169338 | ......G . ....... | Mexico | B | K103R |
| **009** | - | - | KF026125 | G...... . ....... | Ethiopia | C | K101R |
| **016** | - | - | DQ878845 | .C..... . ....... | Italy | B | K101H |
| **017** | - | - | DQ518437 | ......G . ......A | Bahamas | B | K103R, V106I |
| **018** | - | - | JN671242 | ..C.... . .G..... | Argentina | B | K102Q, K104R |
| **019** | - | - | AM181805 | .G..... . ......A | Burkina Faso | CRF06_cpx | V106I |

**Table S3. Comparison of probe design approaches to ensure high coverage.**

| Approach | Methodology | Sequences Used | K65 Probe-Binding Alleles | Advantages | Disadvantages |
|---|---|---|---|---|---|
| **Subtype Agnostic Consensus Nucleotide** | 1. Calculation of nucleotide frequency at each position.<br>2. Degenerate bases selected >5% frequency. | All available sequences are used to determine the consensus sequence. | ATAAARA-RAARGAY | Allows quick determination of nucleotide positions display high diversity. | Bias arises from subtypes over-represented in the sequence database. |
| **Equally-Weighted Consensus Nucleotide** | 1. Calculation of nucleotide frequency at each position.<br>2. Degenerate bases selected >5% frequency. | Uses an equivalent number of sequences from each subtype. | ATAAARA-RAARGAY | Normalizes for bias due to over- or under-represented subtypes in the sequence dataset. | Gives undue significance to variability in low prevalent subtypes such as circulating recombinant forms. |
| **Equally-Weighted Alleles** | 1. Selection of equal number of sequences from each subtype.<br>2. Determine unique probe-binding site alleles.<br>3. Calculate frequency for each unique allele. | Uses an equivalent number of sequences from each subtype. | ATAAAGA-AAAAGAC [40.5%]<br>ATAAAGA-AAAGGAC [18.8%]<br>ATAAAGA-AAAAGAT [17.6%]<br>ATAAAAA-GAAGGAC [13.9%]<br>ATAAAGA-AAAGGAT [3.8%]<br>ATAAAAA-GAAAGAC [2.4%] | Normalizes for bias due to over- or under-represented subtypes in the sequence dataset. | Overcalls the prevalence of the most common allele due to it being the most prevalent allele in subtypes D, F, and G, which have low patient prevalence. |
| **Prevalence-Weighted Alleles** | 1. Calculate frequency for each unique allele per subtype.<br>2. Adjust allele frequency based on subtype prevalence.<br>3. Combine weighted allele frequencies. | All available sequences are used to determine unique probe-binding site alleles per subtype. | ATAAAAA-GAAGGAC [44.4%]<br>ATAAAGA-AAAAGAC [21.0%]<br>ATAAAGA-AAAAGAT [11.4%]<br>ATAAAGA-AAAGGAC [9.1%]<br>ATAAAAA-GAAAGAC [6.5%]<br>ATAAAAA-GAAGGAT [2.8%] | Increases the coverage of oligonucleotide matches in patients with unknown subtypes. The most prevalent probe-binding site here is most common in subtype C and non-A-to-G subtypes. | Some bias may occur due to HIV-1C being the most prevalent subtype. |

"-" indicates the position of the DRM-discriminating SNP

**Table S4. Database HIV-1 subtype distribution compared to patient population prevalence.**

| Subtype | Prevalence | Database Distribution |
|---|---|---|
| **A** | 9.2% | 5.8% |
| **01_AE** | 4.4% | 5.8% |
| **02_AG** | 7.3% | 2.7% |
| **B** | 10.8% | 55.9% |
| **C** | 50.5% | 16.8% |
| **D** | 4.4% | 3.2% |
| **F** | 0.6% | 0.9% |
| **G** | 4.7% | 1.0% |
| **Other** | 8.0% | 7.9% |

From https://www.hiv.lanl.gov/components/sequence/HIV/geo/misc/HIV-1/all/db_world_data.html

**Table S5. Comparison of standard consensus primers to PANDAA primers.**

A single mismatch was present in both the forward and reverse consensus primers: a C:T mismatch toward the 5' end of the forward primer, and a G:A mismatch at the penultimate 3' position of the reverse primer. Both positions were degenerate bases in the PANDAA primers.

| | Forward (5'–3') | Reverse (3'–5') |
|---|---|---|
| **Template 001** | TACCACACCCAGCAGGGTTAA | TGACAGTACTGGATGTGGGGGATGCATATT |
| **Consensus Primers** | .......T............. | .A............................ |
| **PANDAA Primers** | ....N..Y..H..R..DY... | .R.....RY.R.....R..D........Y. |

Median Cq values (interquartile ranges) of six replicates using either consensus or PANDAA primers across a dynamic range of 001 template DNA.

| Copy Number | Consensus | PANDAA |
|---|---|---|
| **$10^6$** | 21.2 (IQR, 21.2 to 21.3) | 24.7 (IQR, 24.6 to 24.9) |
| **$10^5$** | 25.6 (IQR, 25.5 to 25.6) | 28.2 (IQR, 28.2 to 28.3) |
| **$10^4$** | 28.9 (IQR, 28.7 to 28.9) | 32.0 (IQR, 31.9 to 32.0) |
| **$10^3$** | 36.0 (IQR, 35.6 to 36.3) | 35.6 (IQR, 35.5 to 35.8) |
| **$10^2$** | 42.5 (IQR, 41.3 to 42.8) | 39.3 (IQR, 39.2 to 39.3) |

**Table S6. PDR degeneracy at positions with 1–5% nucleotides present.**

| | 2830F-Forward PANDAA Primer | | | | 2896R-Reverse PANDAA Primer | |
|---|---|---|---|---|---|---|
| **Consensus** | **95%** | **96–97%** | **98%** | **99%** | **95%** | **99%** |
| **Degeneracy** | 288 | 1,536 | 2,048 | 18,432 | 384 | 1,536 |

We varied the PDR to represent the 95–99% consensus (i.e., 1–5% nucleotides present).The forward PANDAA primer, 2830F, had four PDR variations based on nucleotide frequency: 95%, 96/97% (no difference in degeneracy), 98%, and 99%. The reverse primer, 2896R, had only two PDR variations: 95% and 99%. The lowest combined degeneracy with a forward and reverse primer was 672-fold with 2830F+95% + 2896R-95%, and the highest degeneracy was 19,968-fold with the 99% consensus variants.

**Table S7. Effects of PDR degeneracy on PANDAA sensitivity.**

With the 001 template, regardless of PDR degeneracy in the 2896R PANDAA primer, incorporating nucleotides with a frequency of 2–4% (i.e., 96–98% consensus) into the 2830F PANDAA primer had a modest effect, increasing the sensitivity by ~1.25-fold. Including degenerate bases when a nucleotide was present at 1% (i.e., 99% consensus) reduced sensitivity by ~1.08-fold.

| | **2896R–95% PDR Degeneracy** | | | | **2896R–99% PDR Degeneracy** | | | |
|---|---|---|---|---|---|---|---|---|
| **2830F PDR** | **95%** | **96–97%** | **98%** | **99%** | **95%** | **96–97%** | **98%** | **99%** |
| **Median Cq** | 30.02 | 29.68 | 29.69 | 30.12 | 29.83 | 29.51 | 29.48 | 29.93 |
| **St Dev** | 0.09 | 0.03 | 0.14 | 0.10 | 0.03 | 0.08 | 0.10 | 0.15 |
| **ΔCq** | - | -0.33 | -0.33 | 0.11 | - | -0.32 | -0.35 | 0.11 |
| **Fold Change** | - | 1.26 | 1.25 | -1.08 | - | 1.24 | 1.27 | -1.08 |

A greater effect of PDR degeneracy on sensitivity was observed when using template 014, which requires adaptation by both forward and reverse PANDAA primers. Degenerate bases introduced at positions with a nucleotide frequency ≥3% enhanced sensitivity by ~22-fold. A similar trend in increased sensitivity was apparent when using a primer representing the 98% consensus, with a loss in effect occurring when degeneracy matched nucleotides with a frequency of 1%.

| | **2896R-95% PDR Degeneracy** | | | | **2896R-99% PDR Degeneracy** | | | |
|---|---|---|---|---|---|---|---|---|
| **2830F PDR** | **95%** | **96–97%** | **98%** | **99%** | **95%** | **96–97%** | **98%** | **99%** |
| **Median Cq** | 34.93 | 30.98 | 31.29 | 34.69 | 35.02 | 30.55 | 31.33 | 34.91 |
| **St Dev** | 0.04 | 0.10 | 0.05 | 0.06 | 0.45 | 0.15 | 0.04 | 0.03 |
| **ΔCq** | - | -3.96 | -3.64 | -0.25 | - | -4.47 | -3.69 | -0.10 |
| **Fold Change** | - | 15.54 | 12.46 | 1.19 | - | 22.19 | 12.92 | 1.07 |

**Table S8. Effect of a -6G sequential adaptation primer on other templates.**

| Sample Name | dCq | Fold Change | Sequence | Mismatch |
|---|---|---|---|---|
| **AAA 001** | 0.7 | -1.5 | AAAAGAA C AAATCAG | |
| **AAA 002** | 0.3 | -1.2 | ....... . ..G.... | +3G |
| **AAA 003** | -8.6 | 203.2 | .G..... . ....... | -6G |
| **AAA 004** | 0.6 | -1.5 | ....... . .G..... | +2G |
| **AAA 005** | 0.8 | -1.7 | ..C.... . ....... | -5C |
| **AAA 006** | 0.1 | -1.1 | ....... . ......A | +7A |
| **AAA 007** | 1.1 | 0.5 | ...G... . ....... | -4G |
| **AAA 008** | 0.9 | -1.8 | ......G . ....... | -1G |
| **AAA 009** | 0.8 | -1.6 | G...... . ....... | -7G |
| **AAA 010** | 0.6 | -1.4 | ....... . .....T. | +6T |
| **AAA 011** | -2.8 | 5.5 | .G..A.. . ....... | -6G, -3A |
| **AAA 012** | 0.7 | -1.5 | ....... . .....C. | +6C |
| **AAA 013** | 0.9 | -1.7 | ....... . .....G. | +6G |
| **AAA 014** | -0.3 | 1.2 | .G..... . ..G.... | -6G, +3G |
| **AAA 015** | 1.4 | 0.4 | C...... . ....... | -7C |
| **AAA 016** | 5.8 | 0.0 | .C..... . ....... | -6C |
| **AAA 017** | 1.0 | -1.8 | ......G . ......A | -1G, +7A |
| **AAA 018** | 0.8 | -1.6 | ..C.... . .G..... | -5C, +2G |
| **AAA 019** | -2.4 | 4.3 | .G..... . ......A | -6G, +7A |
| **Median** | 0.7 | -1.4 | | |
| **25th percentile** | 0.2 | -1.6 | | |
| **75th percentile** | 0.9 | 0.5 | | |

**Table S9. Individual and total allele-specific pro-amplification primers on PANDAA performance.**

| Template | Probe-Binding Site | Mismatch | Single Pro-Amp dCq | Total Pro-Amp dCq |
|---|---|---|---|---|
| **Probe** | AAAAGAA C AAATCAG | | | |
| **001** | ....... . ....... | | | |
| **002** | ....... . ..G.... | +3G | -0.6 | 0.1 |
| **003** | .G..... . ....... | -6G | -8.3 | -3.2 |
| **004** | ....... . .G..... | +2G | -1.2 | 0.4 |
| **005** | ..C.... . ....... | -5C | 0.0 | -0.5 |
| **006** | ....... . ......A | +7A | -1.3 | 0.1 |
| **007** | ...G... . ....... | -4G | 0.1 | 0.1 |
| **008** | ......G . ....... | -1G | 0.0 | 0.2 |
| **009** | G...... . ....... | -7G | -1.0 | -0.7 |
| **010** | ....... . .....T. | +6T | -0.6 | 0.0 |
| **011** | .G..A.. . ....... | -6G, -3A | -7.2 | -7.6 |
| **012** | ....... . .....C. | +6C | 1.2 | -0.4 |
| **013** | ....... . .....G. | +6G | 0.0 | 0.1 |
| **014** | .G..... . ..G.... | -6G, +3G | -0.4 | -0.1 |
| **015** | C...... . ....... | -7C | -1.1 | -0.5 |
| **016** | .C..... . ....... | -6C | -2.3 | 0.2 |
| **017** | ......G . ......A | -1G, +7A | -1.0 | 0.4 |
| **018** | ..C.... . .G..... | -5C, +2G | -1.0 | 0.2 |
| **019** | .G..... . ......A | -6G, +7A | -1.9 | -1.3 |

Pro-Amplification, Pro-Amp.

**Table S10. Probe-Binding Site Alleles Relative to Wild-Type and DRM Synthetic Templates**

| | K65R * | | | K103N | | | Y181C | | | M184V / I † | | |
|---|---|---|---|---|---|---|---|---|---|---|---|---|
| | 5' ADR | SNP | 3'ADR | 5' ADR | SNP | 3'ADR | 5' ADR | SNP | 3'ADR | 5' ADR | SNP | 3'ADR |
| Probe | **ATAAAGA** | **A/G** | **RAAAGAC** | **AAAAGAA** | **A/C** | **AAATCAG** | **TTATCT** | **A/G** | **YCAATA** | **TCAATAT** | **A/GTR** | **GATGA** |
| | | | | | | | | | | **AATAT** | **ATG/A** | **GATGACT** |
| Allele 001 | .....A. | | ...G... | ....... | | ......A | ...... | | ...... | ....... | | ....... |
| Allele 002 | ....... | | ....... | ....... | | ..G...A | .C.... | | ...... | ......C | | .....T. |
| Allele 003 | ....... | | ......T | .G..... | | ......A | .G.... | | ...... | C...... | | ....... |
| Allele 004 | ....... | | ...G... | ....... | | .G....A | ....T. | | ...... | C.....C | | .....T. |
| Allele 005 | .....A. | | ....... | ..C.... | | ......A | ...... | | ...G.. | ...G... | | .....T. |
| Integrated A | | | | | | | | | | | | |
| 001 | ATAAAAA | A | RAAGGAC | AAAAGAA | A | AAATCAA | TTATCT | A | TCAATA | TCAATAT | ATG | GATGACT |
| 002 | ATAAAGA | A | RAAAGAC | AAAAGAA | A | AAGTCAA | TCATCT | A | TCAATA | TCAATAC | ATG | GATGATT |
| 003 | ATAAAGA | A | RAAAGAT | AGAAGAA | A | AAATCAA | TGATCT | A | CCAATA | CCAATAT | ATG | GATGACT |
| 004 | ATAAAGA | A | RAAGGAC | AAAAGAA | A | AGATCAA | TTATTT | A | CCAATA | CCAATAC | ATG | GATGATT |
| 005 | ATAAAAA | A | RAAAGAC | AACAGAA | A | AAATCAA | TTATCT | A | TCAGTA | TCAGTAT | ATG | GATGATT |
| Integrated B ‡ | | | | | | | | | | | | |
| 001 | ATAAAAA | G | RAAGGAC | AAAAGAA | C | AAATCAA | TTATCT | G | TCAATA | TCAATAT | RTA | GATGACT |
| 002 | ATAAAGA | G | RAAAGAC | AAAAGAA | C | AAGTCAA | TCATCT | G | TCAATA | TCAATAC | RTA | GATGATT |
| 003 | ATAAAGA | G | RAAAGAT | AGAAGAA | C | AAATCAA | TGATCT | G | CCAATA | CCAATAT | RTA | GATGACT |
| 004 | ATAAAGA | G | RAAGGAC | AAAAGAA | C | AGATCAA | TTATTT | G | CCAATA | CCAATAC | RTA | GATGATT |
| 005 | ATAAAAA | G | RAAAGAC | AACAGAA | C | AAATCAA | TTATCT | G | TCAGTA | TCAGTAT | RTA | GATGATT |

* K65 probes were designed against the second most prevalent allele (002) to minimize the number of secondary polymorphisms to be adapted in a single probe-binding site.

† Probes for 184V and 184I differ in the positioning of the discriminating SNP. For 184V, an A → G substitution occurs at the 1st nucleotide of codon 184, and for 184I a G → A substitution at the third nucleotide. This leads to a 2nt downstream shift in the probe-binding site for 184I. As both DRMs are adapted and amplified by the same PANDAA primers the 5' and 3' ADRs must cover the cumulative probe-binding region.

‡ The M184V mutation can represent a change from ATG → GTG or GTA; the M184I mutation is ATG → ATA. To incorporate both 184V and 184I mutations into a single synthetic template, the 184 codon is given as RTA (GTA and ATA) in the Integrated B templates.

**Table S11. Probe-Binding Site Alleles in Patient Samples**

| K65 | Count | |
|---|---|---|
| ATA AAA A-G AAG GAC | 47 | 65.3% |
| ATA AAA A-G AAA GAC | 12 | 16.7% |
| ATA AAG A-A AAA GAC | 8 | 11.1% |
| ATA AAA A-G AAG GAT | 3 | 4.2% |
| ATT AAA A-G AAG GAC | 1 | 1.4% |
| ATA AAA A-G AAG GGC | 1 | 1.4% |

| K103 | Count | |
|---|---|---|
| AA AAG AA- AAA TCA G | 57 | 79.2% |
| AA AAG AA- AAA TCA A | 2 | 2.8% |
| AA AAG AA- AGA TCA G | 2 | 2.8% |
| AG AAG AA- AAA TCA G | 4 | 5.6% |
| AA AAG AA- AAG TCA G | 2 | 2.8% |
| AA CAG AA- AAA TCA G | 3 | 4.2% |
| AA AAG AA- AAG TCA A | 1 | 1.4% |
| AA AAG AA- AAA TCA R | 1 | 1.4% |

| Y181 | Count | |
|---|---|---|
| GTC ATC T-T CAA TAT | 41 | 56.9% |
| GTT ATC T-T CAA TAC | 13 | 18.1% |
| GTT ATC T-T CAA TAT | 8 | 11.1% |
| GTT ATC T-C CAA TAT | 4 | 5.6% |
| GTC ATC T-T CAG TAT | 4 | 5.6% |
| GTC ATC T-T CAA TAC | 1 | 1.4% |
| RTT ATC T-C CAA TAT | 1 | 1.4% |

| M184I | Count | |
|---|---|---|
| AAT --- TGG ATG ACT | 49 | 68.1% |
| AAT --- TGG ATG ATT | 14 | 19.4% |
| AAT --- TAG ATG ACC | 3 | 4.2% |
| AGT --- TGG ATG ACT | 3 | 4.2% |
| AAT --- TGG ATG ACC | 1 | 1.4% |
| AGT --- TGG ATG ATT | 1 | 1.4% |
| AAT --- TAG ATG ACT | 1 | 1.4% |

| M184V | Count | |
|---|---|---|
| TC AAT --- TGG ATG A | 59 | 81.9% |
| CC AAT --- TGG ATG A | 5 | 6.9% |
| TC AAT --- TAG ATG A | 4 | 5.6% |
| TC AGT --- TGG ATG A | 4 | 5.6% |

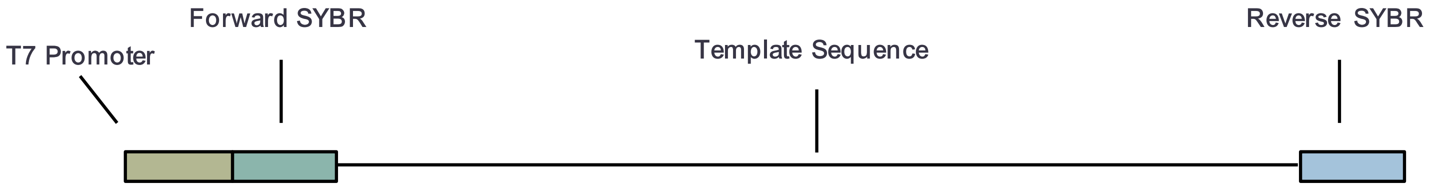

**SYBR primers for quantification:**

```
Fwd 5'-GCT CCT CTG GAA AGG TGA AG
Rev 5'-GCG GAT AAC AAT TTC ACA CAG G
```

**Fig. S1. Template Design.**

Synthetic double-stranded DNA was designed to evaluate the sensitivity and specificity of PANDAA. The 5' region of the template contains the promoter for T7 RNA polymerase to derive synthetic single-stranded RNA. Immediately downstream of the T7 promoter, and at the 3' terminus, we included optimized primer-binding sites to allow SYBR green confirmation of the template copy number. Lyophilized geneStrings (Life Technologies) were resuspended in TE buffer to obtain a template master stock at $10^{10}$ copies/µL. Templates were subsequently diluted in $dH_2O$ supplemented with carrier tRNA from *Saccharomyces cerevisiae* at 0.05 µg/µL (Sigma Aldrich) to provide a dilution series from $10^6$ copies/µL to 100 copies/µL.

#### SYBR qPCR with 2896R-95% Reverse Primer

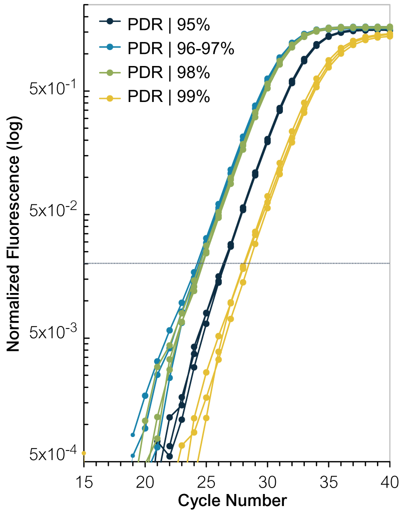

| 2830F PDR | 95% | 96–97% | 98% | 99% |
|---|---|---|---|---|
| **Median Cq** | 26.31 | 24.13 | 24.47 | 28.14 |
| **St Dev** | 0.01 | 0.14 | 0.07 | 0.16 |
| **ΔCq** | - | -2.17 | -1.84 | 1.83 |
| **Fold Change** | - | 4.51 | 3.58 | -3.55 |
| **Amplicon $T_m$** | 73.3°C | 72.8°C | 73.0°C | 72.2°C |

**Melt Curve**

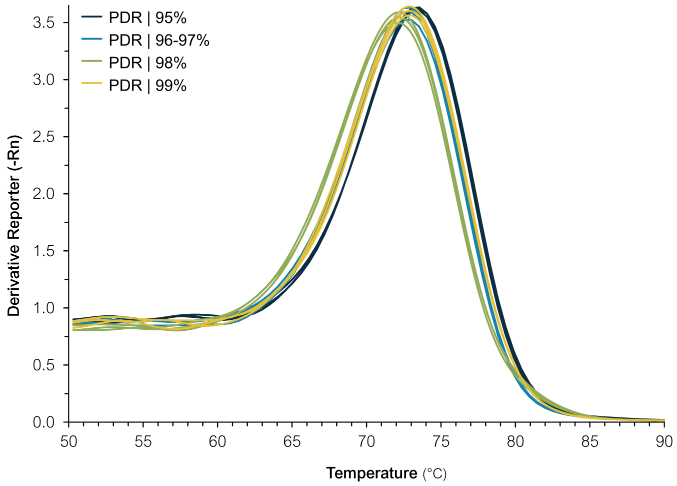

#### SYBR qPCR with 2896R-99% Reverse Primer

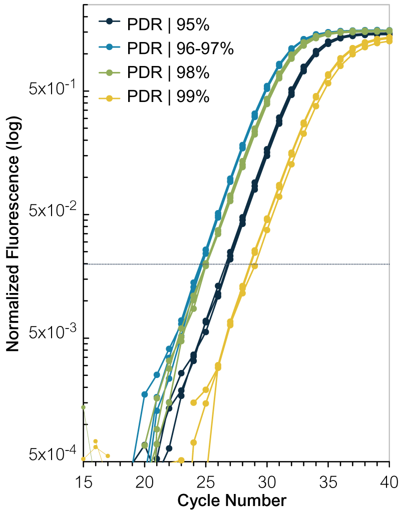

| 2830F PDR | 95% | 96–97% | 98% | 99% |
|---|---|---|---|---|
| **Median Cq** | 26.69 | 24.62 | 24.98 | 28.79 |
| **St Dev** | 0.10 | 0.08 | 0.15 | 0.23 |
| **ΔCq** | - | -1.68 | -1.32 | 2.49 |
| **Fold Change** | - | 3.21 | 2.50 | -5.60 |
| **Amplicon $T_m$** | 72.0°C | 71.4°C | 71.4°C | 71.4°C |

**Melt Curve**

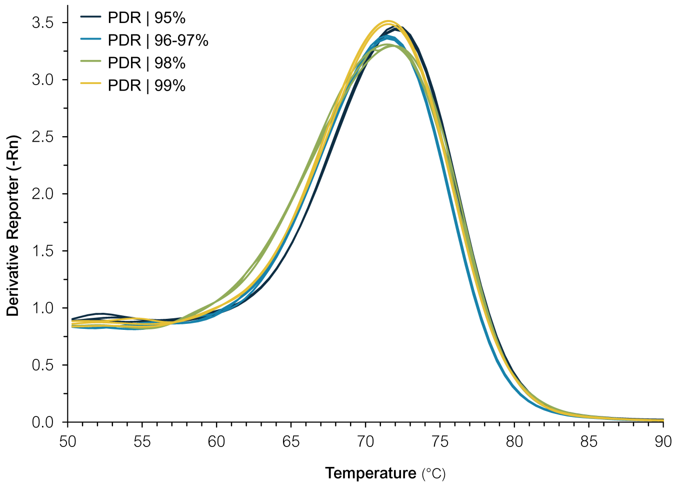

**Fig. S2. Resolution of a single PCR product using a range of PDR degeneracy with SYBR qPCR.**

Each 2830F PDR variant was used with either the 2896R-95% or 2896R-99% in a SYBR green qPCR with the 014 template. A similar pattern of increased reaction efficiency to that with probe-based PANDAA was observed: an increase in degeneracy up to 96–98% consensus improved amplification by both SYBR and probe-based qPCR compared to 95% consensus, then subsequently reduced amplification. This suggests that there is a tipping point after which an increase in PDR degeneracy reduces amplification efficiency. This effect was noticeable only with the forward primer and not the reverse primer because 2830F-99% has a 18,432-fold degeneracy whereas 2896R-99% has only a 1,536-fold degeneracy. The lower amplicon Tm and broader melt curve when using the 2896R-99% primer is indicative of the wider range of GC% content compared to 2896R-95% (25.8 - 58.1% vs. 29.0–54.8%), and therefore the wider range of primer Tm (63.0–78.7°C vs. 64.2–77.3°C).

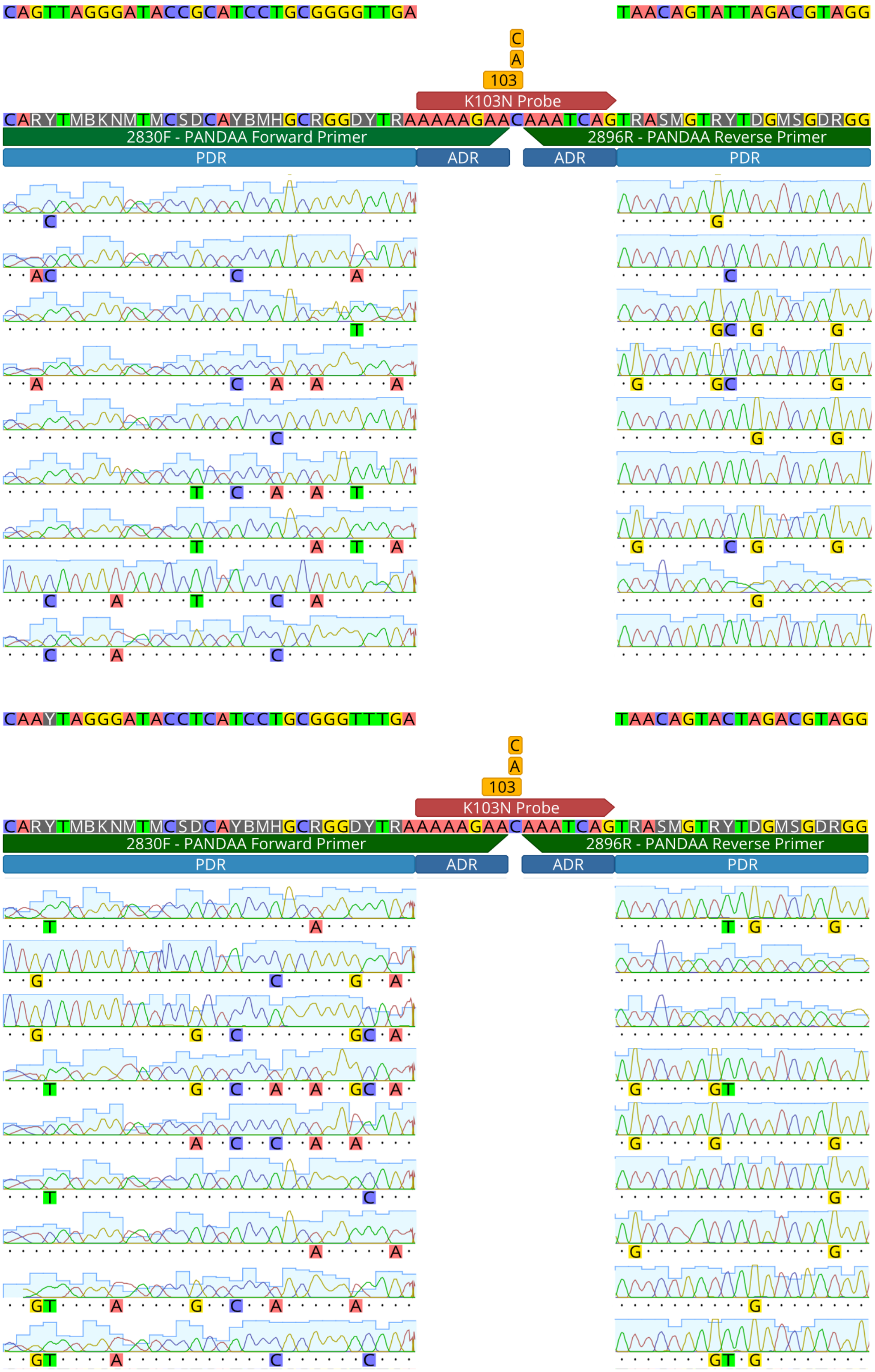

**Fig. S3. Adaptation of the primer-binding region by PANDAA.**

66-bp amplicons, which were derived from PANDAA performed on synthetic DNA templates containing the 001 probe-binding site allele, were cloned into the PCR Cloning Kit (New England Biolabs; Ipswich, MA, USA), and individual clones were sequenced using the associated cloning analysis forward and reverse primers. Representative chromatograms from two DNA templates with different primer-binding sites are shown.

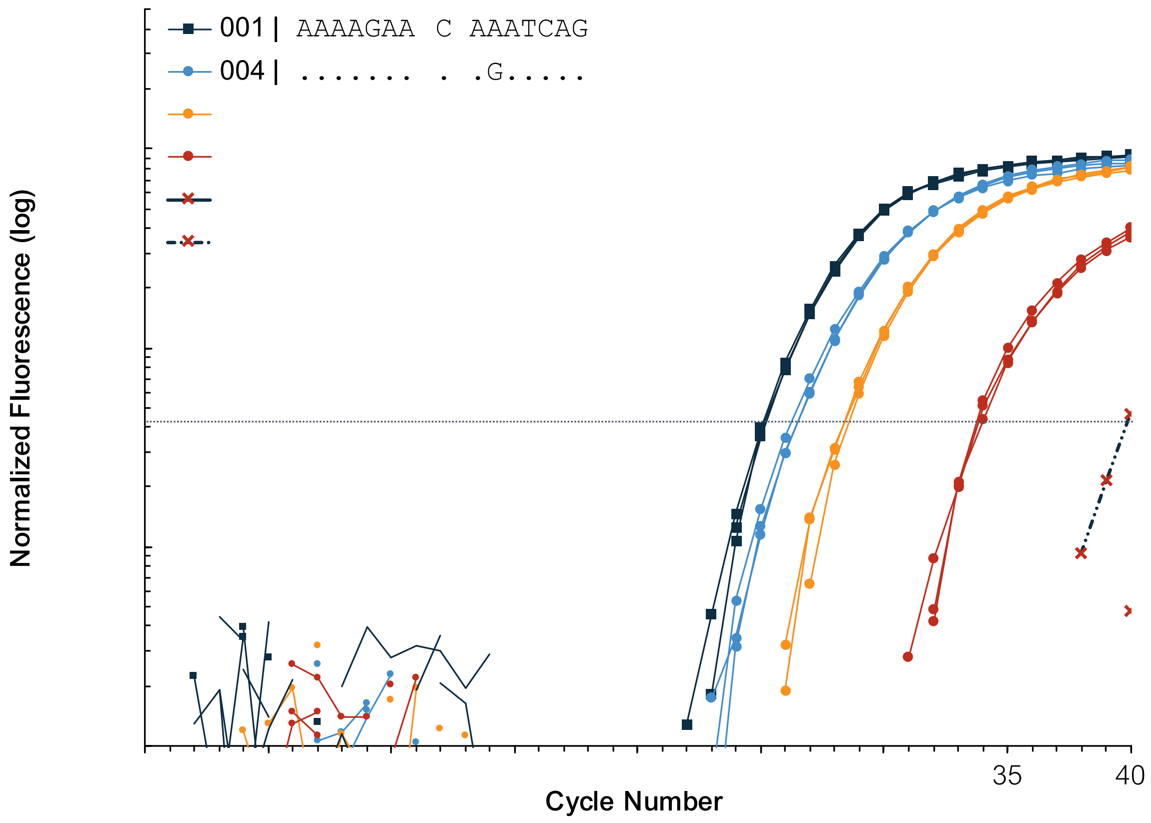

**Fig. S4. PANDAA in a one-step RT-qPCR with RNA.**

Synthetic RNA was derived from synthetic DNA constructs using the HiScribe T7 High Yield RNA Synthesis Kit, NEB. After *in vitro transcription*, DNA was removed using RQ1 RNase-Free DNase and RNA was purified using RNeasy MiniElute Cleanup Kit (Qiagen; Hilden, Germany) with additional on-column DNA digestion. Single-stranded RNA was quantified using the Qubit RNA High Sensitivity assay (Life Technologies; Carlsbad, CA, USA) and diluted in $dH_2O$ supplemented with *S. cerevisiae* carrier tRNA at 50 ng/μL (Sigma Aldrich).

$10^5$ copies of RNA derived from templates 001, 004, 007, and 011 were used in a PANDAA reaction with 15U MMLV RT (NEB) added to Kapa Probe Fast Master Mix in a final volume of 10μL. In a one-step RT-qPCR, the reaction was incubated at 42°C for 15 minutes followed by the standard cycling conditions for the K103 PANDAA. $dH_2O$ with carrier RNA was included in a no RT control to verify no carryover DNA contamination from the RNA synthesis. Templates 004 and 007, which have a probe-binding site mismatch in the 3' and 5' adaptor regions, respectively, demonstrated similar sensitivity in a one-step RT-qPCR. Template 011, which has two mismatches in the 5' adaptor region, was amplified less efficiently, which was also demonstrated using DNA.

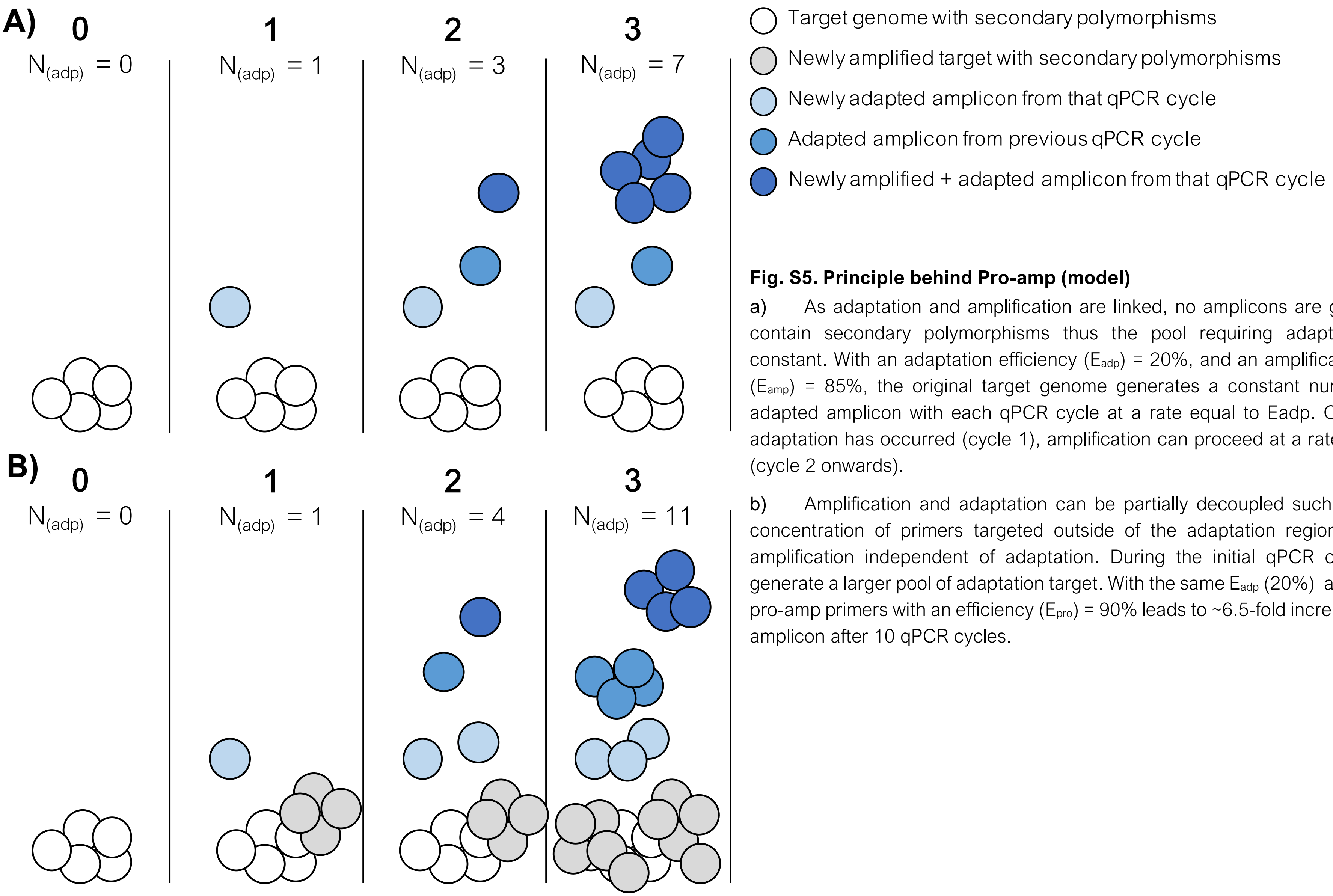

**Fig. S5. Principle behind Pro-amp (model)**

a) As adaptation and amplification are linked, no amplicons are generated that contain secondary polymorphisms thus the pool requiring adaptation remains constant. With an adaptation efficiency ($E_{adp}$) = 20%, and an amplification efficiency ($E_{amp}$) = 85%, the original target genome generates a constant number of newly adapted amplicon with each qPCR cycle at a rate equal to Eadp. Once the initial adaptation has occurred (cycle 1), amplification can proceed at a rate equal to $E_{amp}$ (cycle 2 onwards).

b) Amplification and adaptation can be partially decoupled such that a limited concentration of primers targeted outside of the adaptation region will promote amplification independent of adaptation. During the initial qPCR cycles this will generate a larger pool of adaptation target. With the same $E_{adp}$ (20%) and $E_{amp}$ (85%), pro-amp primers with an efficiency ($E_{pro}$) = 90% leads to ~6.5-fold increase in adapted amplicon after 10 qPCR cycles.
